# Protein-protein interactions with G3BPs drive stress granule condensation and gene expression changes under cellular stress

**DOI:** 10.1101/2024.02.06.579149

**Authors:** José M. Liboy-Lugo, Carla A. Espinoza, Jessica Sheu-Gruttadauria, Jesslyn E. Park, Albert Xu, Ziad Jowhar, Angela L. Gao, José A. Carmona-Negrón, Torsten Wittmann, Natalia Jura, Stephen N. Floor

**Author notes:** Correspondence: J.M.L-L., S.N.F.

## Abstract

Stress granules (SGs) are macromolecular assemblies that form under cellular stress. Formation of these condensates is driven by the condensation of RNA and RNA-binding proteins such as G3BPs. G3BPs condense into SGs following stress-induced translational arrest. Three G3BP paralogs (G3BP1, G3BP2A, and G3BP2B) have been identified in vertebrates. However, the contribution of different G3BP paralogs to stress granule formation and stress-induced gene expression changes is incompletely understood. Here, we identified key residues for G3BP condensation such as V11. This conserved amino acid is required for formation of the G3BP-Caprin-1 complex, hence promoting SG assembly. Total RNA sequencing and ribosome profiling revealed that disruption of G3BP condensation corresponds to changes in mRNA levels and ribosome engagement during the integrated stress response (ISR). Moreover, we found that G3BP2B preferentially condenses and promotes changes in mRNA expression under endoplasmic reticulum (ER) stress. Together, this work suggests that stress granule assembly promotes changes in gene expression under cellular stress, which is differentially regulated by G3BP paralogs.

## INTRODUCTION

Stress granules (SGs) are membrane-less organelles composed of ribonucleoprotein (RNP) complexes that condense into macromolecular assemblies upon cellular stress^1^. Association of untranslated mRNAs and the enrichment of proteins from the translational machinery into SGs has led to the model that these RNP condensates inhibit mRNA expression and translation during the stress response^2,3^. For instance, stress granule assembly leads to suppression of *Csnk2a1* mRNA translation preventing axonal growth post-injury^4,5^. In yeast, Ded1p condensation into stress granules during heat shock promotes the translation of stress-induced transcripts, while suppressing the translation of housekeeping genes^6^. However, it has been reported that SGs are dispensable for stress-induced global translational arrest^7^. Moreover, active translation of mRNA molecules has been identified inside SGs^8^, leading to uncertainty around the function of stress granules in mRNA expression and their relevance during the stress response.

Stress granules assemble by the condensation of RNAs and RNA-binding proteins such as Ras-GTPase-activating protein (SH3 domain)-binding proteins (G3BPs). G3BPs are common components and crucial SG nucleators under multiple cellular stresses. Knockout of G3BPs inhibits the formation of stress granules under oxidative and endoplasmic reticulum (ER) stress^7,9^. Previous work revealed that G3BP1 participates in protein interaction networks that drive SG assembly via liquid-liquid phase separation of core SG proteins^9–11^. Upon stress-induced translational arrest, multivalency with RNA is increased by binding of G3BP1 with proteins such as UBAP2L and Caprin-1, promoting condensation of RNP complexes into SGs. The G3BP1-Caprin-1 interaction is mediated by the nuclear transport factor 2 like domain (NTF2L) of G3BP1 and a short linear motif of Caprin-1^12^. Perturbing this interaction can lead to disruption of stress granule assembly during cellular stress.

Three G3BP paralogs have been identified in vertebrates: *G3BP1*, and the two splice isoforms encoded by *G3BP2*: G3BP2A, and G3BP2B^13^. Aside from SG assembly, G3BPs are involved in RNA metabolism and in the response against viral infection by either promoting or inhibiting the replication of virus^14^. Furthermore, they interact with key cellular pathways during the progression of certain cancers and neurodegenerative diseases^15,16^. The function of G3BP paralogs in SG assembly has been considered predominantly redundant^9^. For this reason, most research has been focused on understanding the function of G3BP1. However, there is growing evidence suggesting that G3BP paralogs may have different roles in certain biological contexts, such as regulation of mTOR signaling and the response against poliovirus infection^17,18^.

G3BPs condense into SGs upon activation of stress response programs such as the integrated stress response (ISR). The ISR is mediated by stress-sensing kinases (PKR, PERK, HRI, and GCN2) that phosphorylate translation initiation factor eIF2⍺ causing global changes in gene expression^19–21^. Previous work showed that G3BP1 condensation differs across multiple ISR-dependent stressors^22^. However, the role of G3BP1 condensation in the regulation of gene expression during the ISR is not well understood. Furthermore, the effects of different ISR-dependent stimuli such as oxidative and ER stress on G3BP2 condensation and function remains unclear.

In this study, we investigated the function of stress granules by identifying G3BP1^V11A^, a mutant that perturbs G3BP1-Caprin-1 interaction in cells, as an inhibitor of G3BP1 condensation during the stress response. To explore the role of SGs in mRNA expression during the stress response, we performed ribosome profiling (Ribo-seq), a technique that measures ribosome density on mRNAs^23^, and total RNA sequencing (RNA-seq). We found that blocking protein-protein interactions that promote G3BP1 condensation led to marginal changes on mRNA levels and translation under oxidative stress via arsenite (NaAs) treatment. To better understand differences between G3BP paralogs, we studied the function of G3BP1/2 condensation into stress granules upon activation of the integrated stress response via oxidative and ER stress. By performing fluorescence microscopy and SG imaging, we found that G3BP paralogs assemble differently into SGs under ER stress. Under NaAs stress, all three G3BP paralogs robustly form SGs with similar properties. In contrast, while G3BP1 condenses poorly under thapsigargin (Tg) treatment, G3BP2B robustly assembles granules in cells. Furthermore, we found that deficiency of protein-protein interactions that drive G3BP2B condensation corresponds to substantial changes in the expression and translation of specific mRNAs under ER stress. Together, we find that G3BP2B potentiates stress granule formation and gene expression changes under ER stress, indicating G3BP paralogs differentially influence gene expression programs under cellular stress.

## MATERIAL and METHODS

### Plasmids

All G3BP paralogs (G3BP1, G3BP2A, and G3BP2B) cDNAs were cloned by Gibson assembly (NEB, E2611L) into either an mEGFP-C1 (Addgene, 54759) or mEGFP-N1 (Addgene, 54767) plasmid with Kanamycin resistance cassettes. Fusion proteins were interspaced with a glycine-serine (GS) linker of 8 residues to improve folding and stability of flanking domains. G3BP1/2 mutants were generated by site directed mutagenesis. For lentivirus expression, these synthetic constructs were cloned by Gibson assembly into a lentivirus backbone with a SFFV promoter, and an ampicillin resistance cassette for bacterial selection. For the generation of cell lines with labeled nuclei, a H2B-mCherry construct cloned in a lentivirus backbone was kindly shared by Dr. Xiaokun Shu (University of California, San Francisco, USA). For *in vitro* reconstitution, NTF2 domain of G3BP1 variants was cloned into a pET28 MBP-TEV bacterial backbone (Addgene, 69929) with Gibson assembly.

### Cell Culture

Cells were grown in Dulbecco’s Modified Eagle Medium/Ham’s F-12 media, supplemented with 10% Fetal Bovine Serum and 1x of a penicillin streptomycin solution (100x, 10,000 I.U. penicillin (per mL), 10,000 µg/mL streptomycin) and kept at 37°C, 5% CO_2_. Cells were tested for mycoplasma and maintained at a low passage number.

### Immunofluorescence

U-2OS wild type and U2OS G3BP1/2 knockout (KO) cells^7^ were seeded at a 40-50% confluency in triplicate in a glass-bottom 96 well plate with #1.5 cover glass (Cellvis, P96-1.5H-N) in Dulbecco’s Modified Eagle Medium/Ham’s F-12 media, supplemented with 10% Fetal Bovine Serum and 1x of a penicillin streptomycin solution (100x, 10,000 I.U. penicillin (per mL), 10,000 µg/mL streptomycin). Cells were incubated overnight at 37°C, 5% CO_2_. Then, they were treated with 100 µL of 200 µM sodium arsenite (LabChem, LC229001) or 1 µM thapsigargin (Millipore Sigma, 586005-1mg) for 2 hrs to induce oxidative and ER stress, respectively. Treated media was removed and cells were washed once with ice cold 1X PBS. Cells were fixed in 4% paraformaldehyde (Electron Microscopy Sciences, 15710) for 10 min at room temperature. Samples were washed three times with ice cold PBS, incubated with 0.1% Triton x-100 at 4°C for 10 min, and followed with three more washes with PBS at 4°C for 5 min. Blocking was performed under 1% bovine serum albumin (BSA) in PBS-T solution for 30 min at 4°C. Primary antibodies for PABP (Abcam, ab21060, 1:1000), G3BP1 (Proteintech, 13057-2-AP, 1:2000), and G3BP2 (Abcam, ab86135, 1:2000) were incubated in 1% BSA PBS-T overnight at 4°C. Then, samples were washed three times with ice cold PBS for 5 min, and incubated with secondary antibody, Goat anti-Rabbit IgG Alexa Fluor 488 nm (Invitrogen, A11008, 1:2000) for 1 hr at room temperature in the dark in 1% BSA PBS-T. Finally, samples were washed three more times in PBS for 5 min in the dark, and stained with 1x Hoechst solution (Bio-Rad, 1351304) at room temperature for 15 min in the dark. Images were acquired by capturing 5 fields of view per well using a 20x objective (Air objective; N.A. 0.75, W.D 1mm), 100 ms of exposure time for 405 nm and 488 nm lasers at 25% power, on a Nikon CSU-W1/SoRa spinning disk confocal microscope.

### Generation of stable cell lines

U-2OS G3BP1/2 KO cells^7^ seeded in a 6-well plate were infected at a 60-70% confluency with lentivirus expressing mEGFP-GS-G3BP variants (G3BP1, G3BP2A, and G3BP2B, G3BP1^V11A^, G3BP2B^V11A^) and co-incubated with 8 µg/mL polybrene (Millipore Sigma, TR-1003-G). Cells were spun down at 1000xg, 30°C for 2 hrs. Then, infected cells were incubated at 37°C, 5% CO_2_ for 72 hrs. Infected cells were sorted twice to achieve mEGFP positive populations.

### Live-cell imaging

U-2OS G3BP1/2 KO^7^ cells at a 60-70% confluency were infected with lentivirus expressing H2B-mCherry plasmids obtained from the Shu lab at UCSF. Positive mCherry single cell populations were sorted. U-2OS H2B-mCherry positive G3BP1/2 KO cells were seeded at a 40-50% confluency in a 96-well plate in Dulbecco’s Modified Eagle Medium/Ham’s F-12 media, supplemented with 10% Fetal Bovine Serum and 1x of a penicillin streptomycin solution (100x, 10,000 I.U. penicillin (per mL), 10,000 µg/mL streptomycin). Cells were incubated overnight at 37°C, 5% CO_2_. Cells were infected with lentivirus expressing mEGFP-GS-G3BP variants and co-incubated with 8 µg/mL polybrene. Four different concentrations of virus (1-5-10-50 µL) were titered per G3BP variant to achieve a similar range of G3BP expression levels between proteins. Cells were spun down at 1000xg, 30°C for 2 hours and incubated at 37°C, 5% CO_2_. After 72 hrs post-infection, ∼5,000 cells were pooled and seeded in a glass-bottom 96 well plate with #1.5 cover glass in triplicates and incubated overnight at 37°C, 5% CO_2_ in no phenol red Dulbecco’s Modified Eagle Medium (Mediatech 17-205-CV) supplemented with 10% Fetal Bovine Serum, 1x of a penicillin streptomycin solution (100x, 10,000 I.U. penicillin (per mL), 10,000 µg/mL streptomycin), autoclaved 25 mM HEPES buffer, and 1x Glutamax (Fisher Scientific, 35-050-061). G3BP condensation was induced with either 200 µM sodium arsenite or 1 µM thapsigargin. To account for G3BP condensation induced by blue light, cells were also treated with DMSO. Images were captured every 30 minutes for 10 hrs at a 20x magnification (Air objective; N.A. 0.75, W.D 1mm), 100 ms of exposure time for 561 nm and 488 nm lasers, on a customized Ti inverted Nikon microscope with a Spectral Applied Research LMM5 laser merge module, and a Borealis modified Yokogawa CSU-X1 spinning disk head, as previously described^24^.

### Ribosome Profiling library construction

Ribo-seq library preparation was performed as previously described^25^. Briefly, cells at 80-90% confluency in a 15 cm dish were incubated with either 200µM sodium arsenite or 1 µM thapsigargin for 2hrs, then treated with 50 µg/mL cycloheximide (CHX) for 1 minute, and harvested. Cells were also harvested after treatment with either water or DMSO, as controls. Then, harvested cells were washed gently with PBS containing 50 µg/mL CHX and lysed in ice-cold lysis buffer (20mM Tris pH 7.4, 150 mM NaCl, 5 mM MgCl2, 1% v/v Triton x-100, 1 mM DTT, 20-25 U/mL TURBO DNase (Thermo Fisher Scientific, AM2238), 100 µg/mL CHX). Cells were triturated with a 26-gauge needle. Lysate was recovered after spinning for 10 min, 20,000xg, at 4°C. Lysates were treated with RNase I (Ambion, 100 U/µL) at room temperature for 45 minutes in slow agitation. Then, treated with SUPERase Inhibitor (Ambion, 20 U/µL) on ice. Monosomes were recovered by size exclusion chromatography (Illustra MicroSpin Columns, S-400 HR, VWR, 95017-619), and footprint RNA fragments were extracted from the flow-through using a Direct-zol kit (Zymo Research). Gel slices of RNA fragments with sizes between 26-34 nt were excised from a 15% polyacrylamide TBE-Urea gel (Thermo Fisher Scientific, EC68855BOX). Eluted RNA was treated with T4 PNK and preadenylated linker was ligated to the 3’ end using T4 RNA ligase 2 truncated KQ (NEB, M0373L). Linker-ligated RNA was reverse transcribed with Protoscript II (NEB, M0368L) for 30 minutes at 50°C. Template RNA was hydrolyzed with 1M NaOH for 20 minutes at 70°C. Gel slices of reverse transcribed cDNA around 105 nt long, were excised from 15% polyacrylamide TBE-Urea gel. Eluted cDNA was circularized with CircLigase II ssDNA ligase (Lucigen, CL9021K) for 1 hr at 60°C. rRNA was depleted with biotinylated oligos^26^. cDNA libraries were amplified with different reverse indexing primers per sample. Libraries were quantified and checked for quality using a Qubit fluorimeter and Bioanalyzer (Agilent) and sequenced on HiSeq 4000 or NovaSeqX sequencing systems.

### RNA sequencing library construction

RNA-seq library preparation was performed by extracting RNA from 25 µL of lysate with a Direct-zol kit. rRNA was depleted with a NEBNext rRNA Depletion Kit (NEB, E7400), and following manufacturer’s instructions. Libraries were prepared with a NEBNext Ultra II Directional RNA Library Prep Kit of Illumina (E7760), and following manufacturer’s instructions. cDNA libraries were amplified with multiplex oligos for Illumina sequencing (NEB, E6609S). Libraries were quantified and checked for quality using a Qubit fluorimeter and Bioanalyzer (Agilent) and sequenced on HiSeq 4000 or NovaSeqX sequencing systems (single end, 65 nt reads and 100 nt reads, respectively).

### Western blot

U-2OS cells, at a 70-80% confluency, were harvested by scraping from a 6-well plate. Cells were lysed and rotated in RIPA buffer (20 mM Tris-HCl at pH 7.5-7.7, 150 mM NaCl, 1% NP-40, 0.5% DOC, 0.05% SDS) supplemented with a protease inhibitor cocktail (Sigma-Aldrich, 4693159001) for 30 min at 4°C. Samples were analyzed by SDS-PAGE electrophoresis. G3BP1 (Bethyl, A302-033A, 1:1000), G3BP2 (Abcam, ab86135, 1:1000), eIF2⍺ (Cell Signaling Technology, 9722S, 1:1000), p-eIF2⍺ (Abcam, ab32157, 1:1000), GFP (Thermo Scientific, A-11122, 1:1000), Caprin-1 (Proteintech, 15112-1-AP, 1:1000), β-actin (Abcam, ab184092, 1:5000) were detected by western blot. Secondary antibody (LI-COR Biosciences, 926-32211, 1:10000) was detected with a LI-COR Odyssey DLx imaging system.

### Co-Immunoprecipitation of G3BP1-Caprin-1 complex

U-2OS cells, at a 70-80% confluency, were treated with 200 µM NaAs for 2 hours and harvested by scraping from a 10 cm plate. Cells were lysed in lysis buffer (20 mM Tris-HCl pH 7.7, 150 mM NaCl, 5 mM MgCl_2_, 1 mM DTT, 0.5% NP-40, 10% glycerol) containing 1 U/µL RNAse I and a protease inhibitor cocktail. Cells were rotated for 30 minutes at 4°C, and the lysate was cleared by centrifugation at 5000 rpm for 5 minutes at 4°C. GFP-trap agarose beads (ChromoTek, gta-10) were equilibrated in ice-cold wash/dilution buffer (20 mM Tris-HCl pH 7.7, 150 mM NaCl, 5 mM MgCl_2_). Lysates were rotated end-over-end with the equilibrated beads for 2 hours at 4°C. After binding, beads were washed with a wash/dilution buffer twice. Beads were then resuspended in SDS buffer (5% β-mercaptoethanol, 30% Glycerol, 10% SDS, 250 mM Tris-Cl, pH 6.8) and analyzed by SDS-PAGE electrophoresis and western blot.

### Protein Expression and Purification

BL21 (DE3) *E.coli* bacteria were transformed with NTF2L domain of G3BP1 variants and cultivated in agar plates at 37 °C overnight. Colonies were selected by kanamycin resistance and expanded in a 1 L Terrific Broth culture media, supplemented with a potassium phosphate buffer (17 mM KH_2_PO_4_,72 mM K_2_HPO_4_) and incubated at 37°C while shaking at 200 rpm. Once the culture reached an OD between 0.7-0.9, 1 mM IPTG was added to induce the expression of the MBP-NTF2L proteins at 18°C overnight. Bacterial pellets were collected by centrifugation for 30 min at 4°C and 4,500 rpm. Pellets were resuspended in 40 mL lysis buffer (50 mM sodium phosphate pH 8.0, 250 mM sodium chloride, 30 mM imidazole, 20 µg/mL DNase I (Roche, #10104159001), 20 µg/mL RNase A (QIAGEN, #19101), 1 tablet EDTA-free protease inhibitor (Roche, #11836170001). After pellets were fully resuspended, they were further lysed by sonication (30 sec on, 30 sec off, 25% amplitude). The sonicated lysates were then spun down for 30 min at 4°C and 40,000 rpm. The supernatant was filtered with 0.45 µm filters before loading onto a His-Trap HP column (Cytiva, #17524801) connected to an AKTA HPLC purification system. The column was equilibrated with buffer A (20 mM sodium phosphate at pH 8.0, 300 mM sodium chloride, 30 mM imidazole) and protein was eluted with a linear gradient of buffer B (20 mM sodium phosphate at pH 8.0, 300 mM sodium chloride, 500 mM imidazole). Eluted protein was cleaved with 1 mg TEV protease and 5 mM β-mercaptoethanol at 4°C overnight to remove histidine tags. Cleaved protein was purified by collecting the flowthrough on a His-Trap column. Then, collected protein was purified by size exclusion chromatography (SEC) with a Superdex 75 Increase 10/300 GL column (Cytiva, #29148721) equilibrated with 10 mM Tris-Cl pH 8.0, 150 mM NaCl. Finally, MBP tags that co-eluted with NTF2L proteins were purified by incubating with amylose resin (New England BioLabs, #E8021L) overnight at 4°C and collecting the flow through the following day. Collected protein was concentrated to 1 mg/mL using a 3kDa concentrator, flash-frozen, and stored at –80°C.

### Differential Scanning Fluorimetry

Differential scanning fluorimetry was performed in a 96-well clear PCR plate (Bio-Rad, #MLL9601) with 2 µM recombinant NTF2L domain, 5x SYPRO orange (Sigma, #S5692) and filled to 20 µL with SEC buffer (10 mM Tris-Cl pH 8.0, 150 mM NaCl). A control well included SEC buffer alone with 5x SYPRO orange. The PCR plate was sealed and loaded to a RT-PCR (Bio-Rad, #1855201) machine where the plate was incubated at 25°C for 1 minute followed by gradual temperature increases by 0.2°C increments until the instrument reached 95°C. Samples were excited at 535 nm and emission collected at 559 nm. The melting temperature (T_m_) was calculated by taking the first derivative of the melting curve.

## DATA ANALYSIS

### Stress granule segmentation

Image analysis was performed by developing CellProfiler pipelines^27^. Briefly, for immunofluorescence (IF) and live-cell imaging experiments, both nuclear and cytoplasmic signals were rescaled by dividing pixel intensities by a factor. Nuclei were segmented with the Otsu thresholding method. Single cells were segmented by propagation from the nuclear signal. PABP and G3BP foci were segmented by enhancing their speckle-like feature and applying either a manual or the Otsu thresholding method for IF and live-cell imaging experiments, respectively. Finally, potential segmentation artifacts were filtered based on eccentricity, where foci less than 0.875 were kept (Fig. S1). Further data processing and plotting was done in python and R.

### Preprocessing and alignment of NGS data

Next-generation sequencing data was processed as previously described^28^. Briefly, Ribo-seq footprint reads were trimmed of adapter sequences with cutadapt and filtered based on a 32 nt cutoff. UMI sequences were collapsed and removed. Reads were aligned with bowtie2 2.4.1 and filtered against a collection of repeat RNAs including tRNA, rRNA, snRNA, srpRNA, among others (RepeatMasker). Filtered reads were mapped to the hg38 version of the human genome using STAR^29^ 2.7.5a with –-sjdbOverhang set to 29. Quality control of mapped footprint reads was performed with R package Ribo-seQC^30^ and MultiQC tool^31^. Spearman correlations were calculated with ggstatsplot^32^ R package. RNA-seq and Ribo-seq profiles had spearman correlation factors of 0.8-1.0 per condition, suggesting good reproducibility among replicates (Fig. S5A-B, S11A-B, S14A-B). As expected, the frequency of fragment lengths for ribosome footprints showed peaks around ∼28 nt, and three nucleotide periodicities at open reading frames (ORFs) of transcripts^23^ (Fig. S5C-D, S11C-D, S14C-D). Deletions at both exons 2 of endogenous *G3BP1* and *G3BP2* genes^7^ to create functional G3BP1/2 knockouts were validated by RNA-seq and Ribo-seq (Fig. S5E-F).

### Differential expression analysis

Log base 2-fold change (LFC) of differentially expressed genes was estimated with DESeq2^33^. A ‘local’ fit for both total RNA and ribosome footprint reads was applied in DESeq2. The *lfcShrink* function was applied to estimate more accurately LFC values of genes with low counts^34^. Only genes with total RNA baseMean and ribosome occupancy baseMean above 20 were considered. Genes with RNA LFC more or less than 0 and p adjusted values less than 0.01 were identified as *Buffering up* or *Buffering down*, respectively. Genes with Ribo LFC more or less than 0 and p adjusted values less than 0.05 were identified as *Ribo ocp up* or *Ribo ocp down*, respectively. Genes changing significantly (p adjusted less than 0.05) at both RNA and ribosome occupancy were identified as either *RNA abundance up* or *RNA abundance down*. Translation efficiency (TE) was estimated with Riborex package^35^. Genes with TE LFC more or less than 0, and p adjusted values less than 0.05, were identified as *TE up* or *TE down*, respectively.

### TE and RNA correlations

Correlations of ΔTE and ΔRNA were plotted with R package ggplot2. Adjusted R^2^ coefficient was calculated with *stat_poly_eq* function and regression line was estimated with *stat_poly_line* function from the ggpmisc package.

### Gene set enrichment analysis

Gene lists generated by DESeq2 were sorted in decreased order based on their RNA-seq LFC scores. GSEA was performed with *gseGO* function from clusterProfiler R package and genome wide annotation for human “*org.Hs.eg.db”*. P values were adjusted with the “fdr” method and a cutoff of 0.05. GSEA was performed with 10000 permutations and ontology defined by biological processes. For SG-associated mRNAs GSEA, gene lists generated by DESeq2 were sorted in decreased order based on their Ribo-seq LFC scores and analyzed with the *fgsea* function.

### Gene ontology analysis

Background was defined by determining all genes with transcripts per million (TPMs) above 1 in the RNA-seq read count datasets. GO was performed with the *enrichGO* function from clusterProfiler R package and genome wide annotation for human “*org.Hs.eg.db”*. P values were adjusted with the “BH” method and a cutoff of 0.05.

## RESULTS

### Key residues in the G3BP1 NTF2L domain are necessary for G3BP1 condensation

To better understand the role of G3BP granules during the stress response, we sought to block protein-protein interactions that lead to SG assembly to study the effect of SG deficiency in cells. Previous work has shown that knockout of endogenous G3BP1/2 inhibits the formation of stress granules under oxidative and ER stress^7,9^. We validated this via IF by measuring condensation of PABP, a common SG protein, in wild type and G3BP1/2 KO cells under NaAs and Tg treatments, which cause oxidative and ER stress^1,36,37^, respectively. As expected, we observed a significant decrease in the percentage of cells with PABP foci under both NaAs and Tg (Fig. S2). By performing live-cell imaging on G3BP1/2 KO cells expressing transgenic G3BP1 N-terminally tagged to monomeric EGFP and treated with 200 µM NaAs, we confirmed previous reports that truncating the intrinsic disordered regions, IDR2 and IDR3, disrupts G3BP1 condensation^9–11^ (Fig. S3B-E). On the other hand, mutating the S149 residue located at the IDR1 of G3BP1 into phosphomimetic and phospho-incompetent amino acids did not decrease the condensation of G3BP1 under oxidative stress (Fig. S3B-E), as previously reported^38^.

G3BP1 IDRs are involved in multiple functions such as RNA-binding^9^. Therefore, the truncation of these domains can impair important functions of G3BPs outside of their role in SG formation, complicating the understanding of the role of SGs in mRNA expression. Instead, we sought to perturb protein-protein interactions driving G3BP1 condensation by mutagenesis of amino acid residues in the NTF2L domain of G3BP1. The NTF2L domain is implicated in mediating interactions between G3BPs and other scaffold proteins, such as Caprin-1 and UBAP2L^7,9,10,12^. In fact, the NTF2L domain’s role in G3BP condensation is exploited by viruses which outcompete interactions between G3BPs and host proteins to prevent stress granule assembly and improve viral replication^39–41^. We therefore generated twelve variants of G3BP1 including alanine substitutions at conserved amino acids (Fig. 1A-B). Within the NTF2L domain, these mutations were proximal to the G3BP1-Caprin-1 complex interface^12^ (Fig. S3F), and therefore potentially implicated in regulating this interaction during stress granule assembly.

**Figure 1:**
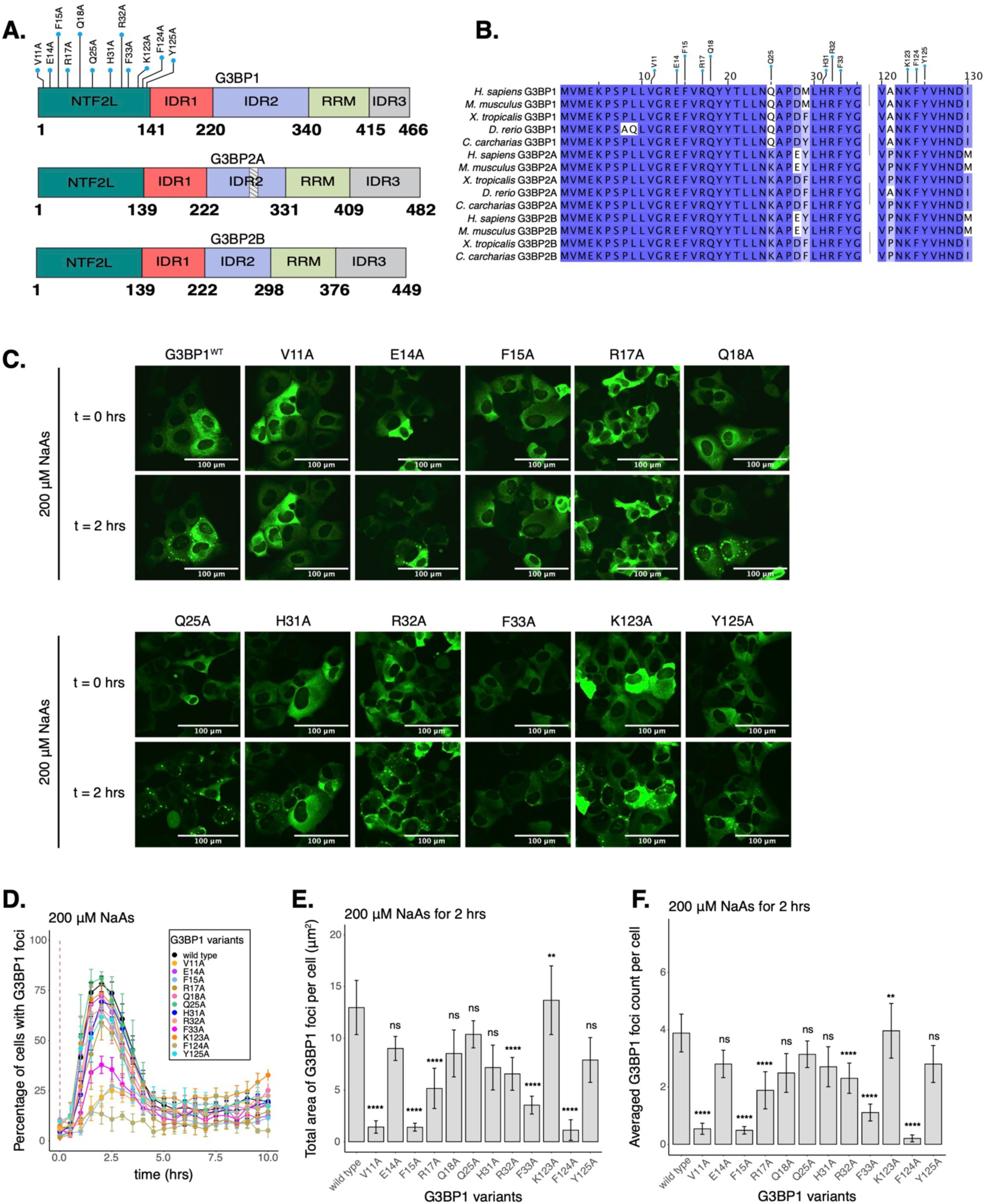
Select residues in the NTF2L domain are key for stress granule formation. **A**. Schematic of G3BP1 (top) domains showing the location of NTF2L alanine substitutions. Schematics of G3BP2A (middle) and G3BP2B (bottom) are also shown. Striped box on G3BP2A shows the region that is spliced out of the G3BP2B IDR2. **B**. Conservation of indicated G3BP NTF2L-domain regions across vertebrate species. **C**. Images (20X objective) of cells expressing mEGFP-G3BP1 variants at t = 0 hr and t = 2 hrs post-treatment with 200 µM NaAs. Puncta in the t = 2 hrs timepoint are stress granules. **D**. Percentage of cells with G3BP1 foci. Vertical red dashed line shows when NaAs was added to cells. **E**. Total area of G3BP1 foci per cell at 2 hours under NaAs. **F**. G3BP1 foci count per cell at 2 hours under NaAs. Plots **D-F** are showing mean ± SEM across N_replicates_ ≥ 3. P-values were calculated based on whole cell populations (n_cells_ ≥ 100 per replicate) relative to G3BP1^WT^. * p < 0.05, ** p < 0.01, *** p < 0.001, **** p < 0.0001.

To test the impact of NTF2L domain mutations on G3BP1 condensation, we used live-cell microscopy (Fig. 1C). We found that alanine substitutions of hydrophobic residues V11, F15, F33, or F124 caused a 50-80% decrease in the percentage of cells with G3BP1 foci at 2 hours of 200 µM sodium arsenite treatment (Fig. 1D). Foci area and count per cell were significantly decreased for these mutations (Fig. 1E-F), suggesting that these hydrophobic interactions play a crucial role in G3BP1 condensation. Mutations on polar residues R17 and R32 also significantly decreased G3BP1 condensation, suggesting that they may mediate important molecular interactions, while most mutated polar residues such as E14, Q18, Q25, and Y125 did not significantly impact the assembly of G3BP1 granules under these conditions (Fig. 1D-F). Interestingly, K123A mutation caused a mild increase in G3BP1 foci area and count per cell (Fig. 1E-F). Prior work identified a mild effect of H31A^12^, which we also observed in our data (Fig. 1D-F). Expression of G3BP1 mutants did not deviate beyond 25% from the median expression of G3BP1^WT^, suggesting that changes in granule properties were not primarily due to differences in protein levels (Fig. S3G). Foci disassembled after 2 hours of stress exposure, confirming the transient nature of G3BP1 condensates under acute stress (Fig. 1D & S3C).

### Mutation of V11 in G3BP1 inhibits association with Caprin-1

To narrow down condensation-deficient mutants to study the function of SGs on mRNA expression, we decided to characterize several SG-deficient mutants *in vitro* (Fig. 2 & S4). The NTF2L domain promotes SG assembly by forming a network with proteins such as Caprin-1 and UBAP2L, which increases the valency of interactions with RNA to drive RNP condensation upon translation arrest and RNA influx^9,10^. Recently, the complex between G3BP1 NTF2L domain and a Caprin-1 short linear motif was characterized as a critical mechanism for SG assembly^12^, giving us a better mechanistic understanding of this interaction. Most of the mutations we generated are located at the interface of the G3BP1-Caprin-1 complex (Fig. S3F) and are predicted to inhibit this interaction. However, substitution of hydrophobic amino acids found in this interface may also destabilize the entire NTF2L domain. To test this possibility, we expressed and purified recombinant NTF2L domain mutants and characterized protein stability (Fig. S4). Purification of G3BP1^F15A^ and G3BP1^F124A^ resulted in low yields due to low protein expression in contrast to G3BP1^WT^ (Fig. S4A-C). This suggests that the F15A and F124A mutants may destabilize the NTF2L domain, thereby leading to their decreased condensation in cells. However, G3BP1^V11A^ and G3BP1^R32A^ expressed robustly and resulted in high protein yields. To measure the stability of these mutants more directly, we performed differential scanning fluorimetry (DSF). Results revealed that G3BP1^V11A^ and G3BP1^R32A^, a milder condensation-deficient mutant, do not significantly decrease the stability of G3BP1 in solution (Fig. 2B-C). To validate the inhibition of the G3BP1-Caprin-1 complex by G3BP1^V11A^, we performed a co-immunoprecipitation assay in cells treated with 200 µM NaAs for 2 hours (Fig. 2D-E). Indeed, we observed a significant decrease in this complex with G3BP1^V11A^, suggesting that granule deficiency is caused by the loss of Caprin-1 interaction. Since we found G3BP1^V11A^ to be a stable variant that cannot interact with Caprin-1, we decided to study the function of this mutant on mRNA expression and translation to better understand the effect of stress granule deficiency during cellular stress.

**Figure 2:**
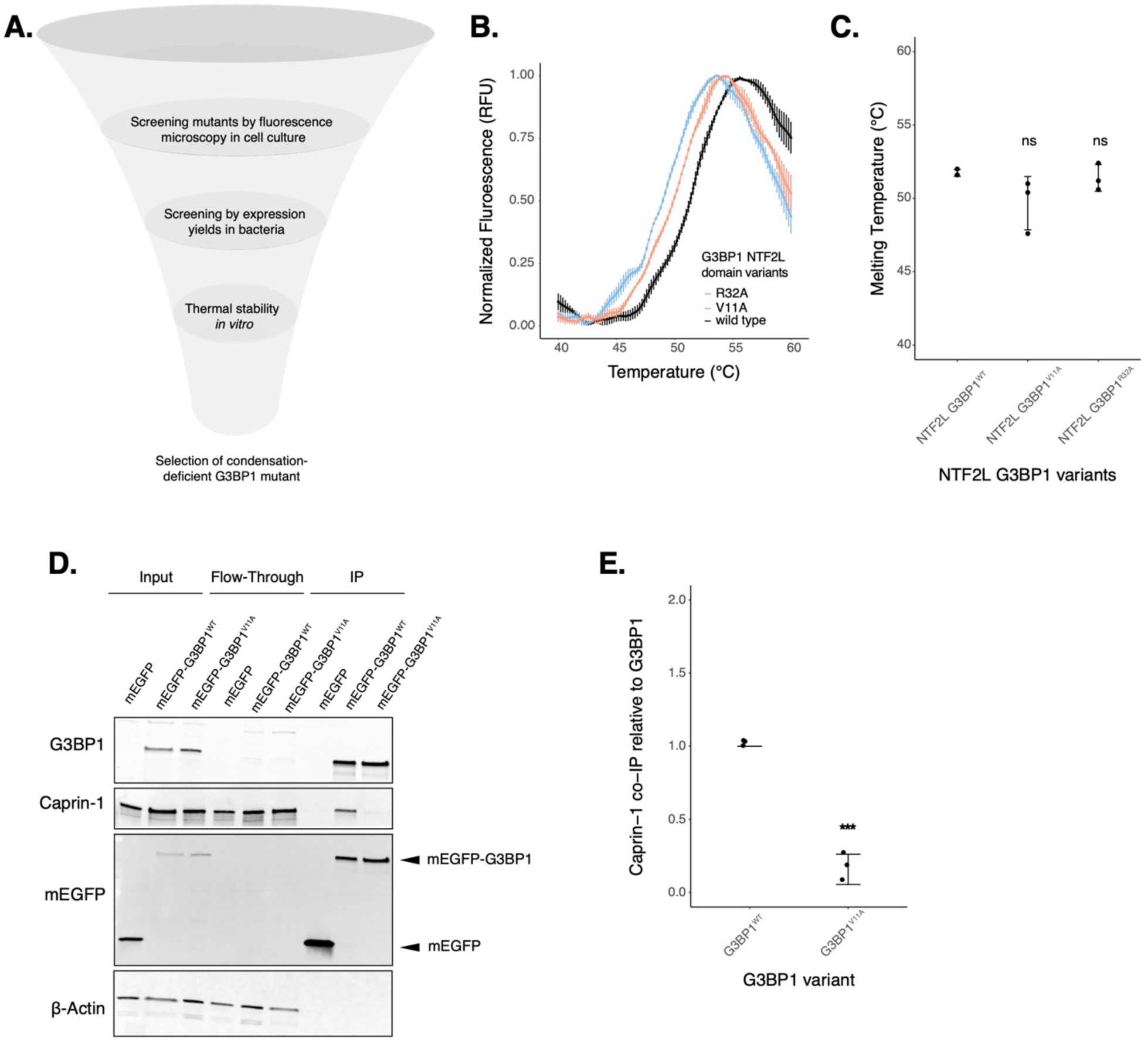
G3BP1V11A inhibits the G3BP1-Caprin-1 complex without affecting protein stability in solution. **A**. Diagram showing screening of G3BP1 mutants (image designed in *biorender.com*). **B**. DSF melting curves for purified G3BP1^WT^, G3BP1^V11A^ and G3BP1^R32A^ NTF2L domains. **C**. Calculated melting temperatures from curves in panel B. **D**. Western blot showing co-immunoprecipitation of the G3BP1-Caprin-1 complex in cells treated with 200 µM NaAs for 2 hours. **E**. Estimations of Caprin-1 band intensities from western on panel D by normalizing on G3BP1 levels. Plots **B,C,E** are showing mean ± SD across N_replicates_ = 3. * p < 0.05, ** p < 0.01, *** p < 0.001, **** p < 0.0001.

### Protein-protein interactions driving G3BP1condensation marginally impact mRNA levels and translation under oxidative stress

The role of stress granules during the integrated stress response is not completely understood. To test the model that G3BP condensation leads to changes in gene expression, we measured global translation using Ribo-seq and total RNA sequencing in U-2OS G3BP1/2 KO cells expressing either transgenic G3BP1^WT^ or G3BP1^V11A^ under the ISR via oxidative stress (Fig. 3). We first confirmed G3BP1 was expressed at comparable levels relative to wild type cells by western blot (Fig. S6A-B). Then, we determined peak phosphorylation of eIF2⍺ occurred around 1-2 hours under oxidative stress (Fig. S6C-D), and therefore harvested cells treated with 200 µM NaAs for two hours. No difference in eIF2⍺ phosphorylation was observed for cells expressing G3BP1^WT^ and G3BP1^V11A^ (Fig. 3B-C). Furthermore, no significant difference in the averaged translation efficiency of ISR canonical factors was observed between G3BP1 variants (Fig. S6E-F), suggesting that stress granule deficiency does not affect ISR activation, as previously reported^7^.

**Figure 3:**
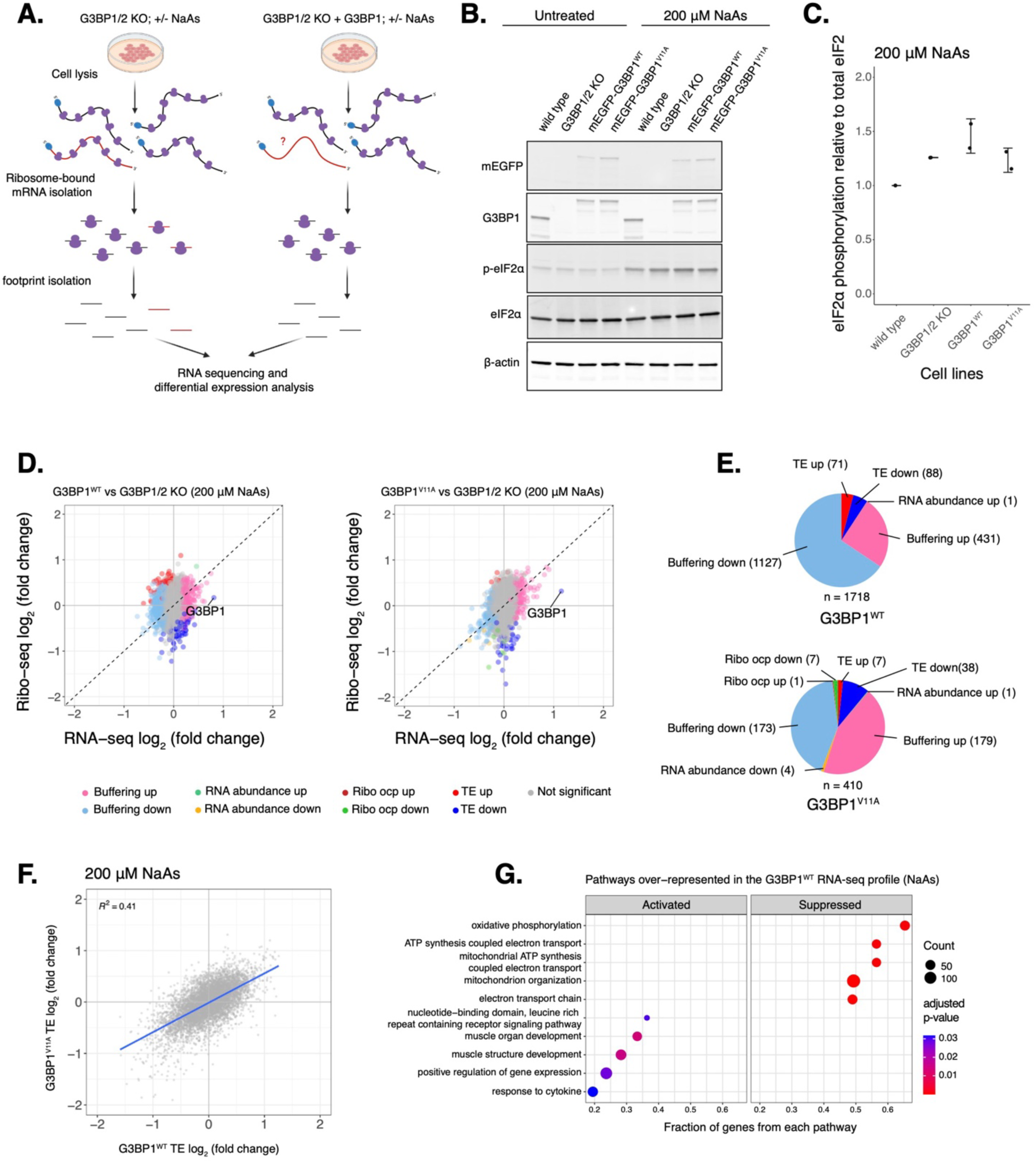
Deficiency in G3BP1 condensation correlates to marginal changes in the expression of select mRNAs under arsenite stress. **A**. Diagram showing Ribo-seq workflow (image designed in *biorender.com*). **B**. Western blot showing levels of eIF2α phosphorylation across U-2OS wild type cells, G3BP1/2 KO cells, and G3BP1/2 KO cells expressing either G3BP1^WT^ or G3BP1^V11A^ N-terminally tagged to mEGFP. Cells were treated either with water for control or 200 µM NaAs for 2 hours. **C**. Comparing levels of eIF2α phosphorylation under NaAs treatment across cell lines from western on panel B. mean ± SD across N_replicates_ = 2. These lysates were used for Ribo-seq and total RNA-seq. **D**. Differential expression plots for Ribo-seq and total RNA-seq. Cells expressing either transgenic G3BP1^WT^ or G3BP1^V11A^ were compared to G3BP1/2 KO cells under 200 µM NaAs for 2 hours. **E**. Count of sensitive genes to G3BP1^WT^ or G3BP1^V11A^ identified on data from panel D. **F**. ΔTE LFC correlations between G3BP1^WT^ and G3BP1^V11A^ under NaAs. **G**. GSEA identifying activated and suppressed pathways by G3BP1^WT^ on the differentially expressed gene sets from RNA-seq.

We determined changes in translation of mRNAs by comparing ribosome profiling and RNA-seq of G3BP1^WT^ or G3BP1^V11A^ to G3BP1/2 KO under oxidative stress (Fig. 3D). Genes with a significant Δ Ribo/RNA ratio were called *TE down* or *TE up*, if their translation efficiency decreased or increased, respectively. If changes in translation were due to corresponding changes in relative levels of RNA, genes were categorized *RNA abundance up* or *RNA abundance down*. While the numbers of G3BP1^WT^ *TE up* and *TE down* genes were 10-fold and 2-fold higher than G3BP1^V11A^ (Fig. 3E), the magnitude of differential TE genes in G3BP1^V11A^/G3BP1^WT^ was marginal (Fig. S6G). Correlation of Δ *TE* genes between G3BP1^WT^ and G3BP1^V11A^ profiles were 0.41 (Fig. 3F). To compare the similarities between G3BP1^WT^ and G3BP1^V11A^, we grouped G3BP1^WT^ sensitive *TE up* and *TE down* mRNAs and measured their ribosome density relative to G3BP1^V11A^. Their ribosome density marginally changed in G3BP1^V11A^/G3BP1^WT^ with median Ribo log_2_ fold changes between 0.13 and –0.12 (Fig. S6H). Interestingly, a gene set enrichment analysis (GSEA) with previously identified SG-associated mRNAs^42^ under NaAs showed that they are significantly suppressed among genes with significant ribosomal occupancy changes (Ribo-seq) for both G3BP1^WT^ (*ES* = –0.22, p = 0.03) and G3BP1^V11A^ (*ES* = –0.29, p < 10^−7^) (Fig. S6I). This suggests that G3BP1 condensation marginally impacts translation of SG-associated transcripts under arsenite stress.

G3BPs mediate RNA stability and expression under physiological and disease contexts^14,15,43^. To decouple the role of G3BP1 condensation into SGs and differential mRNA expression, we identified genes that only changed at the level of RNA without significant changes in ribosome density and termed them as *Buffering up* or *Buffering down*, if their relative abundances increased or decreased, respectively. We found that the number of buffered mRNAs sensitive to G3BP1^WT^ were substantially higher than G3BP1^V11A^-sensitive transcripts (Fig. 3E). *Buffering up* or *Buffering down* genes sensitive to G3BP1^WT^ changed marginally in G3BP1^V11A^/G3BP1^WT^ with median RNA log_2_ fold changes between –0.1 and 0.13, respectively (Fig. S6H). Interestingly, GSEA revealed that G3BP1^WT^ suppresses the expression of genes involved in mitochondrial ATP synthesis (Fig. 3G), implying that G3BP1 participates in the regulation of metabolic changes under arsenite treatment. Overall, these results suggest that perturbing protein-protein interactions driving G3BP1 condensation leads to marginal changes in translation and RNA levels of select transcripts under oxidative stress.

### G3BP paralogs condense differently under ER stress

While our work thus far has focused on G3BP1, three paralogs of G3BPs have been identified in mammals^13^. Even though G3BP paralogs are considered redundant in SG assembly, their functions have been found to differ in mTOR signaling regulation and the response against poliovirus infection^17,18^. Furthermore, G3BP paralogs are differentially expressed across tissues and human diseases^13,44,45^. We therefore hypothesized that they may play differential roles in regulating translation during the ISR. To investigate this idea, we generated G3BP1/2 KO cell lines stably expressing G3BP1, G3BP2A, or G3BP2B fused to mEGFP. To test the role of G3BP paralogs under stress, we performed live-cell imaging on G3BP1/2 KO cells expressing transgenic G3BP paralogs N-terminal tagged with monomeric EGFP and treated with 200 µM NaAs to induce oxidative stress (Fig. 4A-D). G3BP paralogs exhibited minimal differences in the percentage of cells with G3BP foci (Fig. 4B) and on the averaged count of foci per cell (Fig. 4D). No significant differences were observed on the total area of G3BP foci per cell (Fig. 4C). To validate our tagging strategy and confirm our results, we also performed live-cell imaging on G3BP1/2 KO cells expressing C-terminally tagged G3BP proteins (Fig. S7). These results suggest that G3BP1/2 paralogs are redundant under oxidative stress. Similar to G3BP1 condensates, G3BP2 foci disassembled after 2 hours of stress exposure, suggesting that G3BP2 SGs are transient in nature.

**Figure 4:**
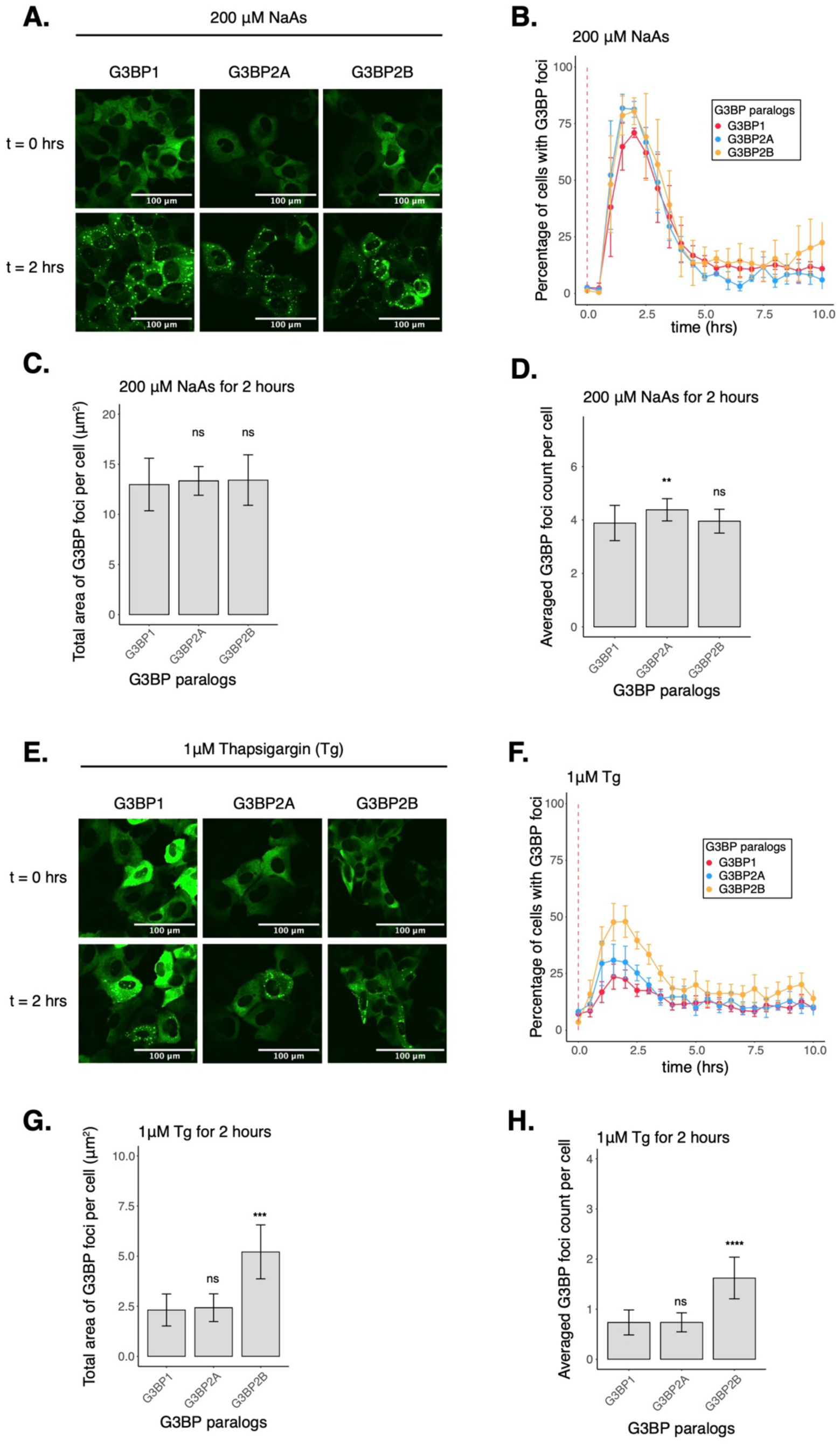
Condensation of G3BP paralogs during the ISR. **A**. Images of cells expressing mEGFP-G3BP paralogs at t = 0 hr and t = 2 hrs post-treatment with 200 µM NaAs. **B**. Percentage of cells with G3BP foci. Vertical red dashed line shows when NaAs was added to cells. **C**. Total area of G3BP foci per cell at 2 hours under NaAs. **D**. G3BP foci count per cell at 2 hours under NaAs. **E**. Images of cells expressing mEGFP-G3BP paralogs at t = 0 hr and t = 2 hrs post-treatment with 1 µM Tg. **F**. Percentage of cells with G3BP foci. Vertical red dashed line shows when Tg was added to cells. **G**. Total area of G3BP foci per cell at 2 hours under Tg. **H**. G3BP foci count per cell at 2 hours under Tg. Plots **B-D** and **F-H** are showing mean ± SEM across N_replicates_ ≥ 3. P-values were calculated based on whole cell populations (n_cells_ ≥ 100 per replicate) relative to G3BP1. * p < 0.05, ** p < 0.01, *** p < 0.001, **** p < 0.0001.

Previous work indicates that properties of G3BP stress granules may differ across stress conditions^22^. Therefore, we also investigated the role of G3BPs in SG assembly under ER stress induced by 1µM Tg (Fig. 4E-H, S8A-D). The percentage of cells with G3BP1 foci under Tg was ∼30% lower than under arsenite stress (Fig. S8B). Stress granule total area and count were also smaller under Tg than arsenite (Fig. S8C-D). Surprisingly, we found that G3BP2B-expressing cells formed more foci under ER stress than G3BP1 or G3BP2A (Fig. 4F). We found that both the total area and count of G3BP2B foci per cell were significantly higher than G3BP1, while no significant differences were observed between G3BP1 and G3BP2A granules (Fig. 4G-H). This suggests that G3BP1 may not be the primary driver of SG formation under ER stress. This also suggests that G3BP2B may form functionally different granules under ER stress, and that the G3BP paralogs differ in their propensity to form SGs under different stressors.

Additionally, expression of G3BP2 isoforms did not deviate beyond 25% from the median expression of G3BP1 (Fig. S8E-F), suggesting that changes in granule properties were not primarily due to differences in protein levels among different cell lines. Interestingly, endogenous G3BP1/2 paralogs in U-2OS wild type cells did not condense differently under either oxidative or ER stress, as revealed by IF (Fig. S9). This may suggest that G3BP2 promotes the condensation of G3BP1 into granules under ER stress by potentially forming G3BP heterodimers in cells, as previously reported^46^.

### G3BP1/2 paralogs regulate mRNA expression and translation differently under ER stress

Given the differences in SG formation across paralogs, we hypothesized that they may play distinct roles in regulating mRNA translation under ER stress. Therefore, we performed Ribo-seq and RNA-seq with G3BP1/2 KO cells stably expressing each transgenic G3BP paralog at comparable levels as revealed by western blot for mEGFP (Fig. S10A-B). G3BP2A/B levels were ∼3-4 times higher than endogenous G3BP2 levels in wild type cells (Fig. S10D). This expression level was selected to match the levels of transgenic G3BP1 in G3BP1/2 KO cells to maintain a constant total amount of G3BP protein, which was similar to the total amount of all G3BP protein in wild-type cells as revealed by Ribo-seq TPMs (Fig. S10I). Phosphorylation of eIF2⍺ under 1 µM Tg peaked around 1-2 hours post-treatment (Fig. S12A-B), therefore cells were harvested at 2 hours for ribosome profiling. ISR activation was validated by the increased expression of ATF4, GADD34, and CHOP (Fig. S12C). Expression of different G3BPs does not affect activation of the ISR, as no significant differences in eIF2⍺ phosphorylation or the average translation efficiency of ISR canonical factors were observed among G3BP paralog cell lines (Fig. 5A-B, S12D).

**Figure 5:**
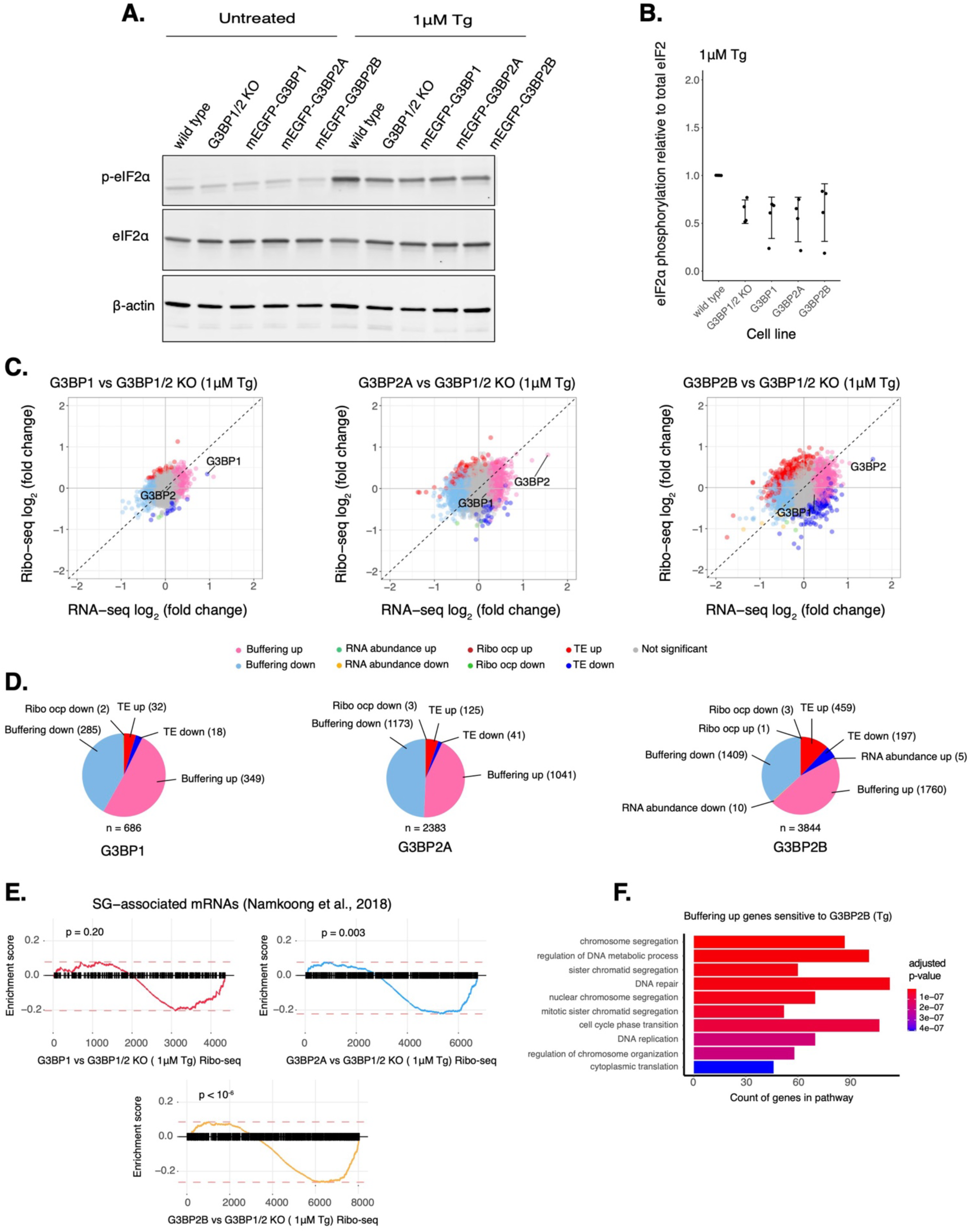
G3BP paralogs impact mRNA expression differently under ER stress. **A**. Western blot showing levels of eIF2α phosphorylation across U-2OS wild type cells, G3BP1/2 KO cells, and G3BP1/2 KO cells expressing either G3BP1, G3BP2A, or G3BP2B N-terminally tagged to mEGFP. Cells were treated either with DMSO for control or 1 µM Tg for 2 hours. **B**. Comparing levels of eIF2α phosphorylation under Tg treatment across cell lines from western on panel A. mean ± SD across N_replicates_ = 4. **C**. Differential expression plots for Ribo-seq and total RNA-seq. Cells expressing either transgenic G3BP1, G3BP2A, or G3BP2B were compared to G3BP1/2 KO cells under 1 µM Tg for 2 hours. **D**. Count of sensitive genes to G3BP1, G3BP2A, or G3BP2B identified on data from panel C. **E**. GSEA for SG-associated mRNAs overlapping with differentially translated gene sets from G3BP1 (upper left), G3BP2A (upper right) and G3BP2B (bottom) Ribo-seq profiles. **F**. GO identifying pathways over-represented in Buffering up genes sensitive to G3BP2B under ER stress.

By comparing each G3BP paralog relative to G3BP1/2 KO cells, we found that the number of differentially translated mRNAs (Δ *RNA abundance* and Δ *TE*) sensitive to G3BP2A/B isoforms was higher than those sensitive to G3BP1, whereas G3BP2B has the biggest impact under ER stress (Fig. 5C-D). Furthermore, there was a higher correlation of Δ *TE* genes between G3BP2A and G3BP2B profiles (0.82), than G3BP1 compared to either G3BP2 isoform (0.53 and 0.43, for 2A and 2B, respectively) (Fig. S13A). This implies that G3BP1/2 paralogs regulate differently the translation of transcripts under ER stress. We also found that buffered mRNA expression (Δ *Buffering*) is different among paralogs where G3BP2 isoforms affected the RNA levels of a higher number of transcripts compared to G3BP1 (Fig. 5D). The correlation of RNA-seq profiles was higher between G3BP2 isoforms (0.85) than G3BP1 compared to either G3BP2A or G3BP2B (0.60 and 0.39, respectively) (Fig. S13B). This implies that G3BP2A/B isoforms also have a greater impact on RNA levels than G3BP1 under ER stress. Moreover, GSEA with previously identified SG-associated mRNAs under Tg^47^ showed that they are suppressed at different magnitudes among genes with significant ribosomal occupancy changes (Ribo-seq) for G3BP1 (*ES* = –0.20, p = 0.20), G3BP2A (*ES* = –0.22, p = 0.003), and G3BP2B (*ES* = –0.26, p < 10^−6^) (Fig. 5E). This result suggests that G3BP paralogs differentially suppress translation of SG-associated transcripts under ER stress.

Gene ontology (GO) analysis revealed that G3BP2B *Buffering up* genes may be involved in multiple pathways such as cell cycle regulation and mRNA translation (Fig. 5F). In fact, previously identified transcripts enriched in granules and known to be involved in cell growth and survival^47^, such as RICTOR, BRCA1, and CREB1, were categorized as *Buffering up* G3BP2B-sensitive mRNAs (Fig. S13C-E). BRCA1 and CREB1 did not significantly changed for either G3BP1 or G3BP2A, however, RICTOR was also identified as a G3BP2A-sensitive *Buffering up* gene. Additionally, dead box helicase DDX3X, an essential gene involved in translation initiation and cell growth^48^, was identified as a *Buffering up* gene sensitive to all G3BP paralogs (Fig. S13F). Interestingly, G3BP1 expression also led to the active expression of genes involved in translation (Fig. S13G). These results highlight both the potential differences and similarities between G3BP paralogs on regulating translation and RNA levels of select transcripts under ER stress.

### Protein-protein interactions driving G3BP2B condensation lead to changes in mRNA expression and translation under ER stress

To determine the potential impact of protein-protein interactions that drive G3BP2B condensation into SGs on gene expression under ER stress, we mutated V11 on G3BP2B by site directed mutagenesis, and performed live-cell imaging on G3BP1/2 KO cells expressing either G3BP2B^WT^ or G3BP2B^V11A^ N-terminally tagged with monomeric EGFP (Fig. 6A). We found that G3BP2B^V11A^-expressing cells exhibited a reduction of the percentage of cells with G3BP2B foci under both oxidative and ER stress (Fig. 6B & Fig. S15B). The total area and count of foci per cell was also significantly decreased (Fig. 6C-D & Fig. S15C-D). This suggests that protein-protein interactions with the V11 residue are critical for the condensation of G3BP2B during the stress response. Expression of G3BP2B^V11A^ mutant did not deviate beyond 25% from the median expression of G3BP2B^WT^ (Fig. S15E), suggesting that changes in granule properties were not primarily due to differences in protein levels.

**Figure 6:**
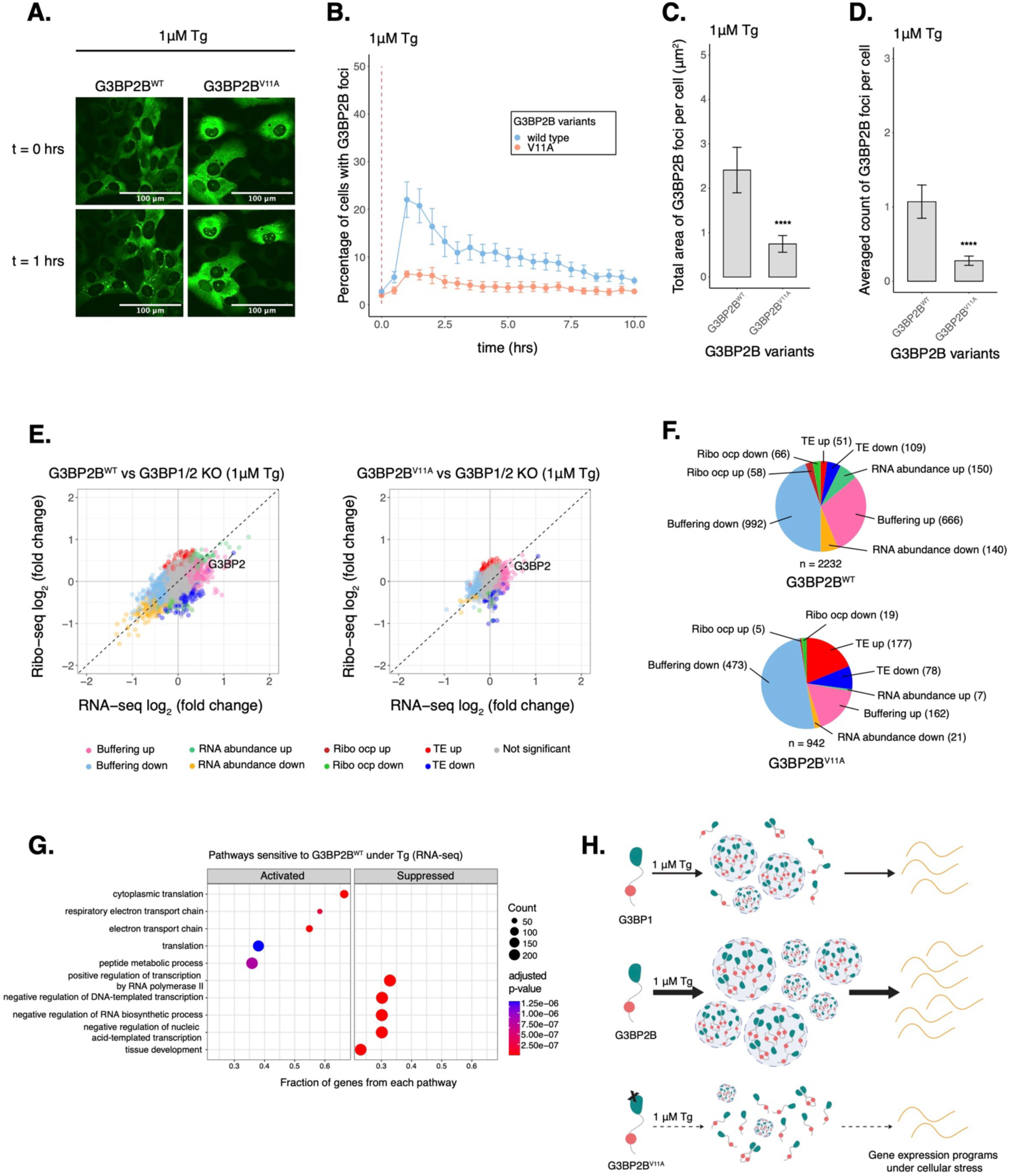
Deficiency in G3BP2B condensation correlates to substantial changes in the expression of select mRNAs under Tg. **A**. Images of cells expressing mEGFP-G3BP2B variants at t = 0 hr and t = 1 hrs post-treatment with 1 µM Tg. **B**. Percentage of cells with G3BP2B foci. Vertical red dashed line shows when Tg was added to cells. **C**. Total area of G3BP2B foci per cell at 1 hour under Tg. **D**. G3BP2B foci count per cell at 1 hour under Tg. Plots **B-D** are showing mean ± SEM across N_replicates_ ≥ 3. P-values were calculated based on whole cell populations (n_cells_ ≥ 100 per replicate). * p < 0.05, ** p < 0.01, *** p < 0.001, **** p < 0.0001. **E**. Differential expression plots for Ribo-seq and total RNA-seq. Cells expressing either transgenic G3BP2B^WT^ or G3BP2B^V11A^ were compared to G3BP1/2 KO cells under 1 µM Tg for 2 hours. **F**. Count of sensitive genes to G3BP2B^WT^ or G3BP2B^V11A^ identified on data from panel E. **G**. GSEA identifying activated and suppressed pathways by G3BP2B^WT^ on the differentially expressed gene sets from RNA-seq. **H**. Model showing the role of protein-protein interactions on the condensation of G3BP1/2 paralogs and mRNA expression during the ISR.

We then performed Ribo-seq and RNA-seq with G3BP1/2 KO cells stably expressing either G3BP2B^WT^ or G3BP2B^V11A^ and harvested after 2 hours of Tg treatment (Fig. 6E). To minimize artifacts caused by higher levels of transgenic G3BP2B, we developed cell lines expressing comparable levels of G3BP2B relative to wild type cells as measured by western blot (Fig. S15F-G). eIF2⍺ phosphorylation did not differ between cells expressing G3BP2B^WT^ and G3BP2B^V11A^ (Fig. S15H) and average translation efficiency of ISR canonical factors was similar across cell lines (Fig. S15I-J).

By comparing both G3BP2B^WT^ and G3BP2B^V11A^ to G3BP1/2 KO cells (Fig. 6E), we found that blocking protein-protein interactions that promote stress granule condensation correlates with decreased abundance of translation repressed transcripts (Δ TE down and Δ RNA abundance down) sensitive to G3BP2B (Fig. 6F). The correlation of Δ *TE* genes between G3BP2B^WT^ and G3BP2B^V11A^ profiles were 0.44 (Fig. S16A). This suggests that G3BP2B condensation represses select mRNAs under ER stress. Moreover, the number of *RNA abundance up* genes was also higher for G3BP2B^WT^, even though the number of *TE up* genes was lower compared to G3BP2B^V11A^. By grouping G3BP2B^WT^-sensitive genes and evaluating their ribosome density in the G3BP2B^V11A^/G3BP2B^WT^ Ribo-seq profile (Fig. S16B), we observed bigger median Ribo log_2_ fold changes in Δ *RNA abundance* (up, –0.222; down, 0.341) compared to Δ *TE* transcripts (up, –0.107; down, 0.121) (Fig. S16C). This may imply that G3BP2B granules impact translation of specific mRNAs by mainly regulating their abundance in bulk cytoplasm. Consistent with these results, we also observed substantial median RNA log_2_ fold changes of both Δ *RNA abundance* and Δ *Buffering* transcripts in the G3BP2B^V11A^/G3BP2B^WT^ RNA-seq profile (Fig. S16C).

We found via GSEA that, under ER stress, genes involved in mRNA translation were activated by G3BP2B^WT^ (Fig. 6G). Furthermore, genes differentially repressed by SG-deficiency (G3BP2B^V11A^/G3BP2B^WT^) are also identified as part of translation via GO analysis (Fig. S16D), suggesting that G3BP2B condensation may improve the expression of specific transcripts involved in cell growth and survival. In fact, DDX3X was identified as a condensation-dependent *RNA abundance down* gene in the G3BP2B^V11A^/G3BP2B^WT^ profile (Fig. S16E), consistent with our previous Ribo-seq experiments. Moreover, GO also revealed that select genes differentially activated by SG-deficiency are members of the *response to wounding* pathway, suggesting a role of G3BP2B condensation on cell migration and the stress response (Fig. S16F). This is consistent with a previous report describing the role of G3BPs on cellular migration^17^. Finally, GSEA also revealed that SG-associated mRNAs identified under ER stress^47^ were translationally suppressed at a higher magnitude by G3BP2B^WT^ (*ES* = –0.31, p < 10^−9^) than G3BP2B^V11A^ (*ES* = –0.24, p < 10^−4^) (Fig. S16G). Overall, these results suggest that perturbing protein-protein interactions driving G3BP2B condensation leads to changes in expression and translation of select mRNAs under ER stress.

## DISCUSSION

To decouple G3BP condensation from other functions such as RNA-binding, we introduced alanine substitutions along the NTF2L domain of G3BP1 that block protein-protein interactions important for G3BP condensation. By performing fluorescence microscopy and SG imaging, we found that hydrophobic residues V11, F15, F33, and F124 have substantial impact on G3BP condensation under arsenite stress (Fig. 1). The NTF2L domain mediates protein-protein interactions with proteins such as USP10, Caprin-1, and UBAP2L^9,10,12^. Macromolecules such as Caprin-1 act as bridges that facilitate phase separation of G3BPs to promote SG assembly^10^. Binding between G3BP1 and Caprin-1 was characterized previously, where the NTF2L domain was shown to interact with Caprin-1 via a short linear motif^12^. NTF2L residues explored in this study are proximal to this interface. Hence, we found that G3BP1^V11A^ mutant inhibits the G3BP1-Caprin-1 complex in cells under NaAs, without affecting protein stability in solution (Fig 2).

The role of stress granules on mRNA translation has remained uncertain. Here we found that perturbing protein-protein interactions that drive G3BP1 condensation corresponds to marginal changes in mRNA expression and translation under NaAs and does not impact levels of ISR activation (Fig 3). The numbers of genes differentially expressed were decreased in cells expressing condensation-deficient G3BP1^V11^, suggesting that SG formation impacts the ability of G3BP1 to regulate expression of select mRNAs during oxidative stress. This also confirms that the ISR is not regulated by G3BP expression or stress granule formation.

In this work, we found that G3BP1/2 paralogs condense similarly under arsenite stress (Fig. 4A-D) while differentially under ER stress induced by thapsigargin treatment. Under Tg stress, G3BP2B condensed in a higher percentage of cells, also leading to bigger granule sizes and numbers per cell compared to either G3BP1 or G3BP2A (Fig. 4E-H). These results may imply that SGs form differently under different stress conditions, as previously reported^22^. Composition and architecture of G3BP1 granules has been previously studied by proteomics, RNA isolation, and super-resolution microscopy techniques^42,49–51^. Our findings motivate performing similar studies focused on G3BP2 granules to better understand the differences and heterogeneity of these condensates from G3BP1 SGs.

Interestingly, we found that granules in U-2OS cells, which naturally express G3BP1, G3BP2A, and G3BP2B, do not condense differently under arsenite versus ER stress (Fig. S9). A previous study showed that G3BP1/2 can form heterodimers in cells^46^. G3BP2, with a higher propensity for condensation due to its increased multivalency^11^, potentially dimerizes with G3BP1 and they cooperate to increase stress granule assembly. Our findings motivate future work to further understand possible cooperative behavior between G3BP paralogs and how this impacts cellular stress response.

Our results also suggest that G3BP1/2 paralogs may have different functions specifically under ER stress. We performed Ribo-seq to test the role of G3BP condensation on mRNA expression during ER stress, and we found that G3BP2B regulates the expression of more genes compared to either G3BP1 or G3BP2A (Fig. 5C-D). These observations may be explained by a higher propensity of G3BP2B for SG-assembly. To test that G3BP2B condensation relates to changes in mRNA expression, we performed Ribo-seq in cells expressing G3BP2B^V11A^, which led to deficiency in G3BP2B condensation under both oxidative and ER stress (Fig. 6A-D & Fig. S15A-D). We found that deficiency of G3BP2B condensation corresponds to a substantial decrease in the number of genes regulated by G3BP2B under ER stress. Furthermore, genes differentially activated in the presence of G3BP2B granules were identified as part of the translational machinery (Fig. 6G & S16D), suggesting the potential role of G3BP condensation on regulating mRNA translation and cell survival during the stress response and recovery. Moreover, we observed that SG-associated transcripts are predominantly translationally repressed by G3BP2B under Tg, supporting the model that SGs downregulate translation of select transcripts via sequestration, particularly under ER stress. However, it is also possible that G3BP2B condensates may be mediating mRNA expression by an alternate mechanism unexplored by this study.

Here, we propose that G3BP condensation into stress granules impact mRNA expression and translation during the ISR. However, there are alternate mechanisms that are potentially playing a direct role on gene expression. Mutating V11 may inhibit interaction networks between G3BPs and proteins critical for mRNA regulation both inside and outside of stress granules. The NTF2L domain mediates dimerization among G3BPs^9–11,41,52,53^. Therefore, this mutation may inhibit dimerization leading to changes in SG assembly and gene expression programs. Our work identifies a relationship between perturbations to protein-protein interactions that are critical for G3BP condensation and gene expression changes under stress. However, more biochemical work is needed to further characterize the mechanism of this G3BP mutant and to better understand the extent of its function on stress granule formation and mRNA regulation.

In conclusion, we identified G3BP^V11A^ to study the function of G3BP condensation. Dysregulation of SGs is correlated to the progression of neurodevelopmental diseases, cancer, and viral infection^14,16,54^. Therefore, mutagenesis of V11 could be utilized to better understand the role of G3BP SGs in these contexts. Finally, we revealed that G3BPs lead to differential mRNA expression, translation, and SG formation, suggesting different roles of G3BP paralogs during the ISR.

## ACKNOWLEDGMENTS

Sequencing was performed at the UCSF CAT, supported by UCSF PBBR, RRP IMIA, and NIH 1S10OD028511-01 grants. Read alignment was performed on the UCSF Wynton High Performance Computing Cluster. Microscopy was partly performed with a next-generation CSU-W1/SoRa spinning disk microscope system funded by NIH S10OD028611-01 grant. Flow cytometry and cell sorting was performed at the UCSF Parnassus Flow Cytometry CoLab. U-2OS G3BP1/2 KO cells were shared by Dr. Nancy Kedersha. Special thanks to all current and former members of the Floor lab, specially Srivats Venkataramanan and Yizhu Lin for providing feedback on RNA sequencing, data analysis, and initial conceptualization of this project. This project was funded by the National Institutes of Health F31GM143845 award (to J.M.L-L.) and R35GM149255 (to S.N.F.). S.N.F. is a Pew Scholar in the Biomedical Sciences, supported by The Pew Charitable Trusts.

*Author contributions*: J.M.L-L. conceptualized, generated reagents, performed experiments and formal data analysis. C.A.E. supported on protein expression and purification. J.S-G., J.E.P and S.N.F. helped on conceptualization. J.S-G. and A.G. provided support on CellProfiler pipeline development. A.X., Z.J., and J.E.P. provided support on ribosome profiling data acquisition, processing, and analysis. J.A.C.N. provided support on DSF data acquisition and analysis. T.W. provided expertise and shared resources for microscopy experiments. N.J. provided expertise and shared resources for *in vitro* experiments. J.M.L-L. and S.N.F. co-wrote manuscript and acquired funding for this project. S.N.F. supervised this project.

## Sequencing data availability

Sequencing data is available at GEO accession numbers GSE254636, GSE254637, and GSE254638.

## Pipelines availability

GitHub: https://github.com/jliboy/SG_segmentation

