## Supplementary Figures for "Protein-protein interactions with G3BPs drive stress granule condensation and gene expression changes under cellular stress"

**Supplemental Figure 1**

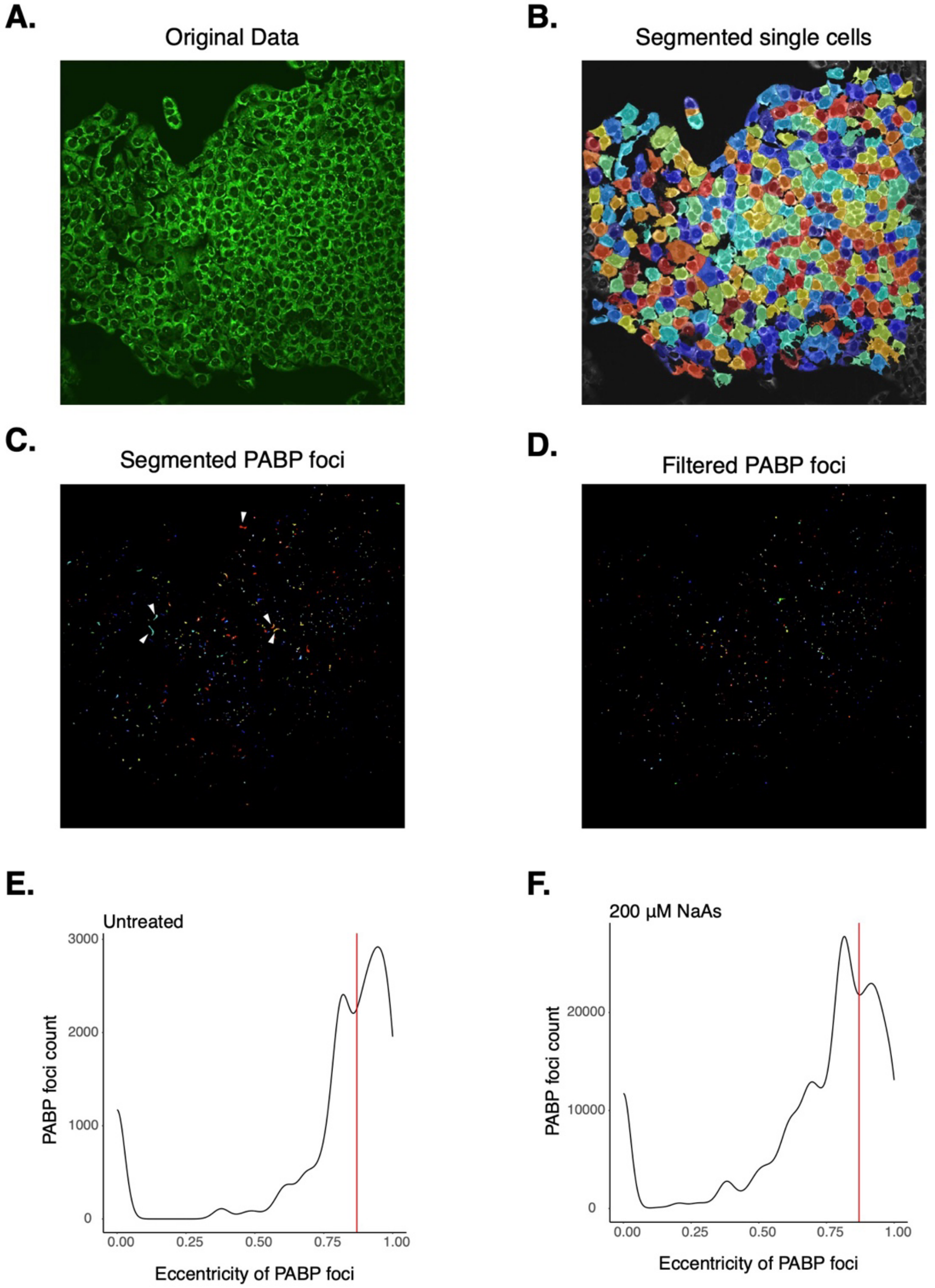


803

**Figure S1: Image analysis pipeline for segmentation of SGs in single cells**. **A**. Image of U-2OS wild type cells forming stress granules under 200 μM NaAs for 2 hours captured by IF. SGs were stained with PABP. **B**. Single cell segmentation. **C**. Segmentation of PABP foci. **D**. Filtered PABP foci based on eccentricity. **E**. Distribution of SG eccentricity under water treatment as a control. SGs above the red vertical line were considered as artifacts. **F**. Distribution of SG eccentricity under 200 μM NaAs for 2 hours. SGs above the red vertical line were considered as artifacts.

#### Supplemental Figure 2

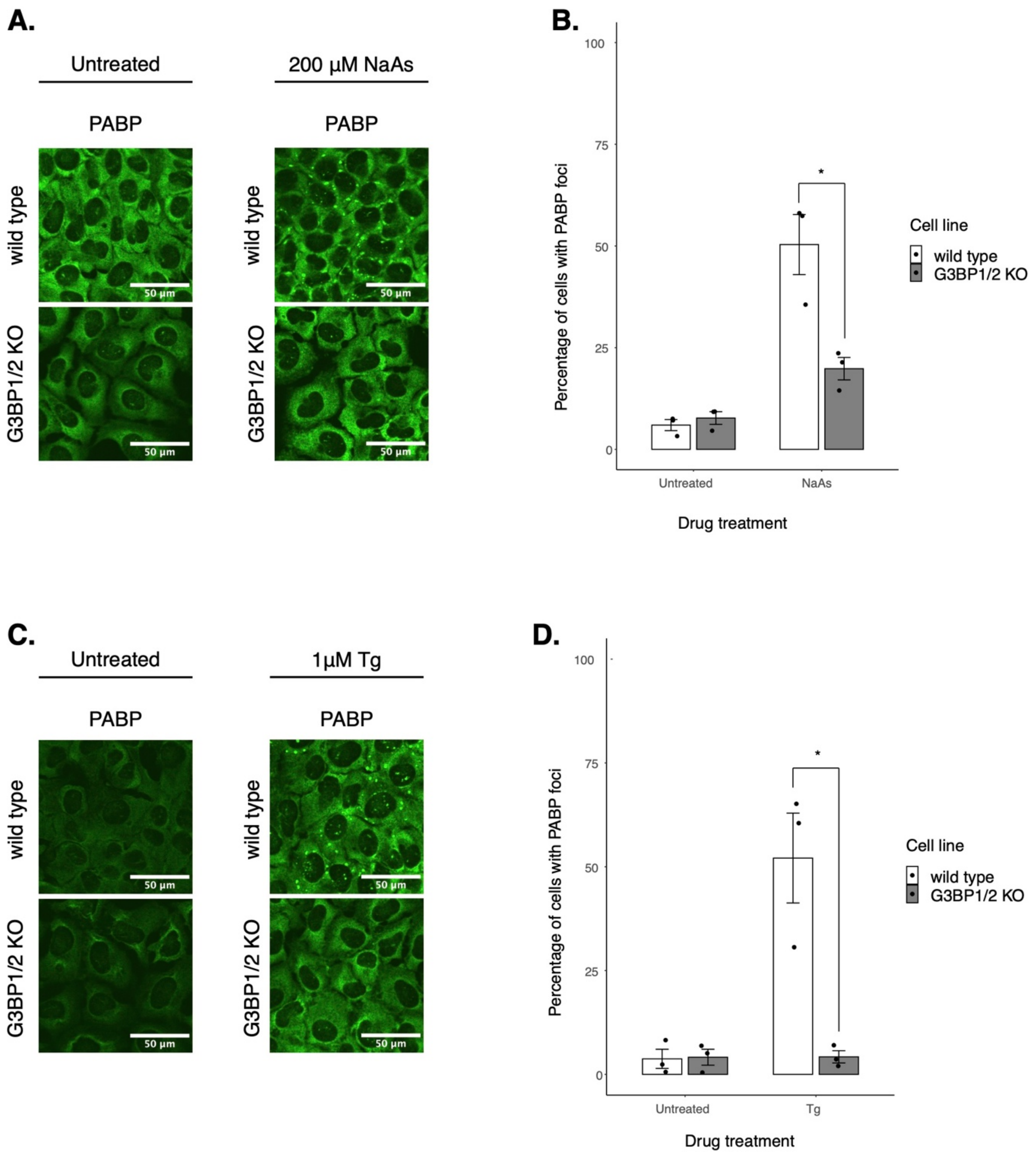

**Figure S2: Inhibition of PABP condensation by G3BP1/2 KO during the ISR**. **A**. SGs stained with PABP by IF. Images are showing U-2OS wild type cells and G3BP1/2 KO cells under water or 200 μM NaAs for 2 hours. **B**. Percentage of cells with PABP foci from data shown in panel A. **C**. SGs stained with PABP by IF. Images are showing U-2OS wild type cells and G3BP1/2 KO cells under DMSO or 1 μM Tg for 2 hours. **D**. Percentage of cells with PABP foci from data shown in panel C. Plots **B** & **D** are showing mean ± SEM across $N_{replicates} = 3$. * $p < 0.05$.

#### Supplemental Figure 3

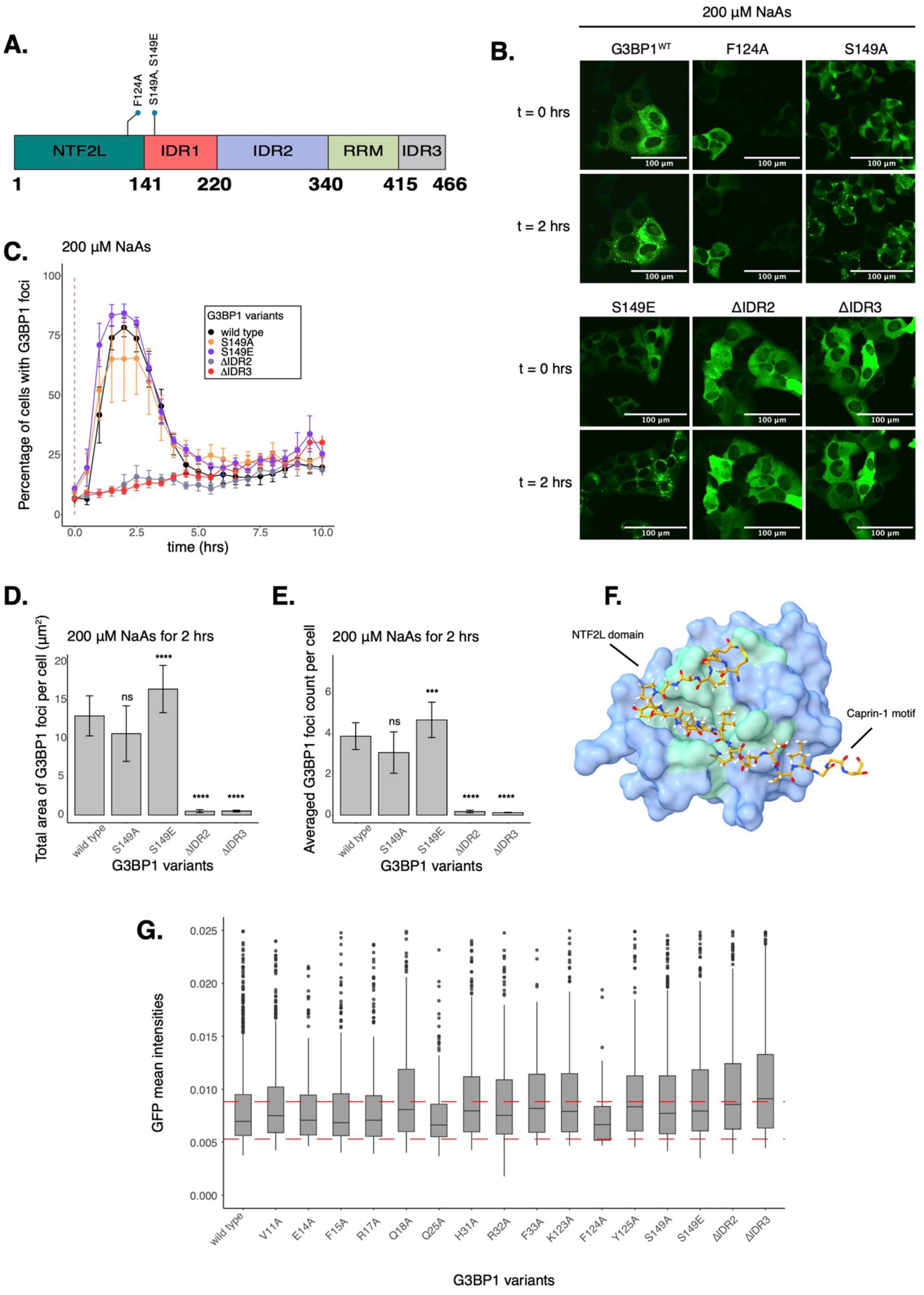


817

**Figure S3: IDRs are critical for G3BP1 condensation under NaAs**. **A**. Schematic of G3BP1 domains showing location of IDRs and S149 residue. **B**. Images of cells expressing mEGFP-G3BP1 variants at t = 0 hr and t = 2 hrs post-treatment with 200 μM NaAs. **C**. Percentage of cells with G3BP1 foci. Vertical red dashed line shows when NaAs was added to cells. **D**. Total area of G3BP1 foci per cell at 2 hours under NaAs. **E**. G3BP1 foci count per cell at 2 hours under NaAs. Plots **C-E** are showing mean ± SEM across $N_{replicates} \geq 3$. P-values were calculated based on whole cell populations ($n_{cells} \geq 100$ per replicate) relative to $G3BP1^{WT}$. * $p < 0.05$, ** $p < 0.01$, *** $p < 0.001$, **** $p < 0.0001$. **F**. Schematic of the G3BP1 NTF2L domain (light blue) interacting with a Caprin-1 motif (gold), PDB ID 6TA7. Location of mutated residues are highlighted (aquamarine). **G**. Cytoplasmic GFP intensities as a proxy for G3BP1 expression across single cells between G3BP1 variants pre-treated with NaAs. Horizontal dashed red lines represent ± 25% from $G3BP1^{WT}$ median cytoplasmic GFP intensity.

#### Supplemental Figure 4

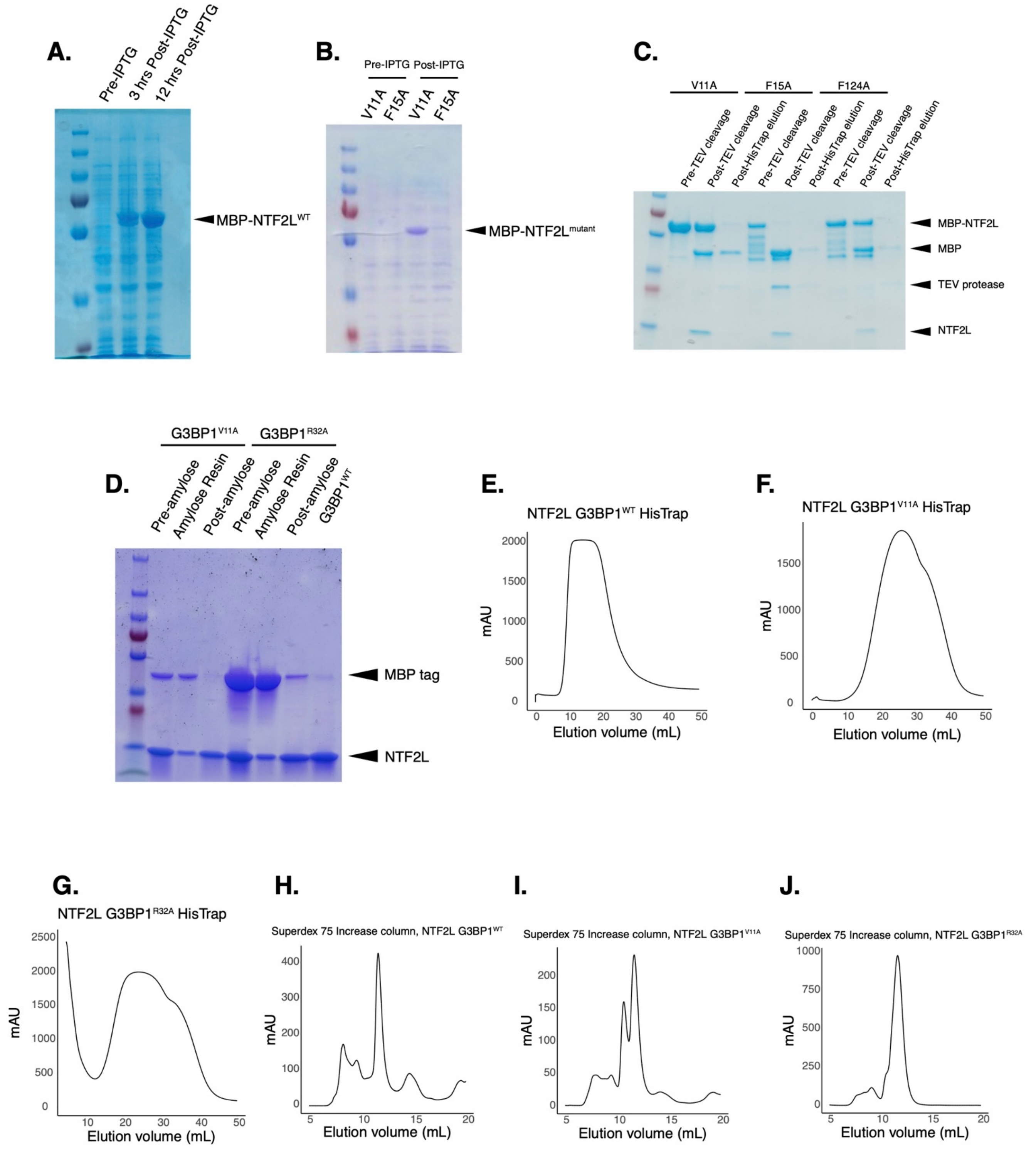

**Figure S4: Expression and purification of recombinant NTF2L proteins**. **A**. Coomassie blue gel showing induced expression of G3BP1$^{WT}$ NTF2L protein. **B**. Coomassie blue gel showing induced expression of G3BP1$^{V11A}$ and G3BP1$^{F15A}$ NTF2L proteins. **C**. Coomassie blue gel showing cleavage of G3BP1$^{V11A}$, G3BP1$^{F15A}$, and G3BP1$^{F124A}$ MBP-NTF2L proteins by TEV protease. **D**. Coomassie blue gel showing Amylose-affinity purification of G3BP1$^{V11A}$ and G3BP1$^{R32A}$ NTF2L proteins after His-Trap and SEC. **E-G**. His-Trap chromatograms for G3BP1$^{WT}$, G3BP1$^{V11A}$ and G3BP1$^{R32A}$ NTF2L proteins. **H-J**. SEC chromatograms for G3BP1$^{WT}$, G3BP1$^{V11A}$ and G3BP1$^{R32A}$ NTF2L proteins.

#### Supplemental Figure 5

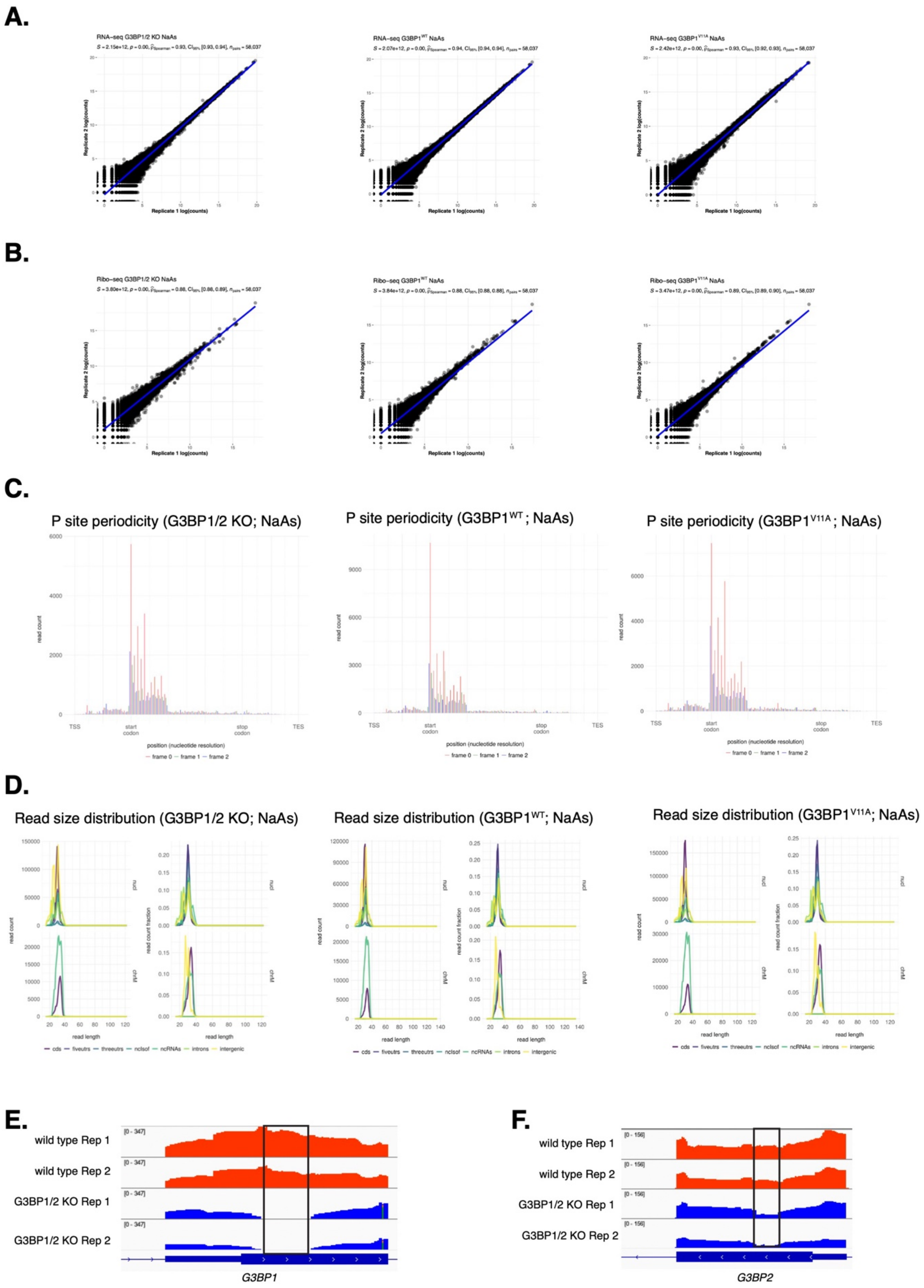


836

**Figure S5: Sequencing QC data for G3BP1/2 KO, G3BP1$^{WT}$, and G3BP1$^{V11A}$ profiles**. **A**. Spearman correlation of G3BP1/2 KO (left), G3BP1$^{WT}$ (middle), and G3BP1$^{V11A}$ (right) RNA-seq sample replicates under NaAs. **B**. Spearman correlation of G3BP1/2 KO (left), G3BP1$^{WT}$ (middle), and G3BP1$^{V11A}$ (right) Ribo-seq sample replicates under NaAs. **C**. P site three nucleotide periodicity for Ribo-seq reads of G3BP1/2 KO (left), G3BP1$^{WT}$ (middle), and G3BP1$^{V11A}$ (right). **D**. Read length distributions of different mRNA species captured by Ribo-seq for G3BP1/2 KO (left), G3BP1$^{WT}$ (middle), and G3BP1$^{V11A}$ (right). **E**. Read coverage tracks showing the site of G3BP1 knockout validated by RNA-seq. **F**. Read coverage tracks showing the site of G3BP2 knockout validated by RNA-seq.

### Supplemental Figure 6

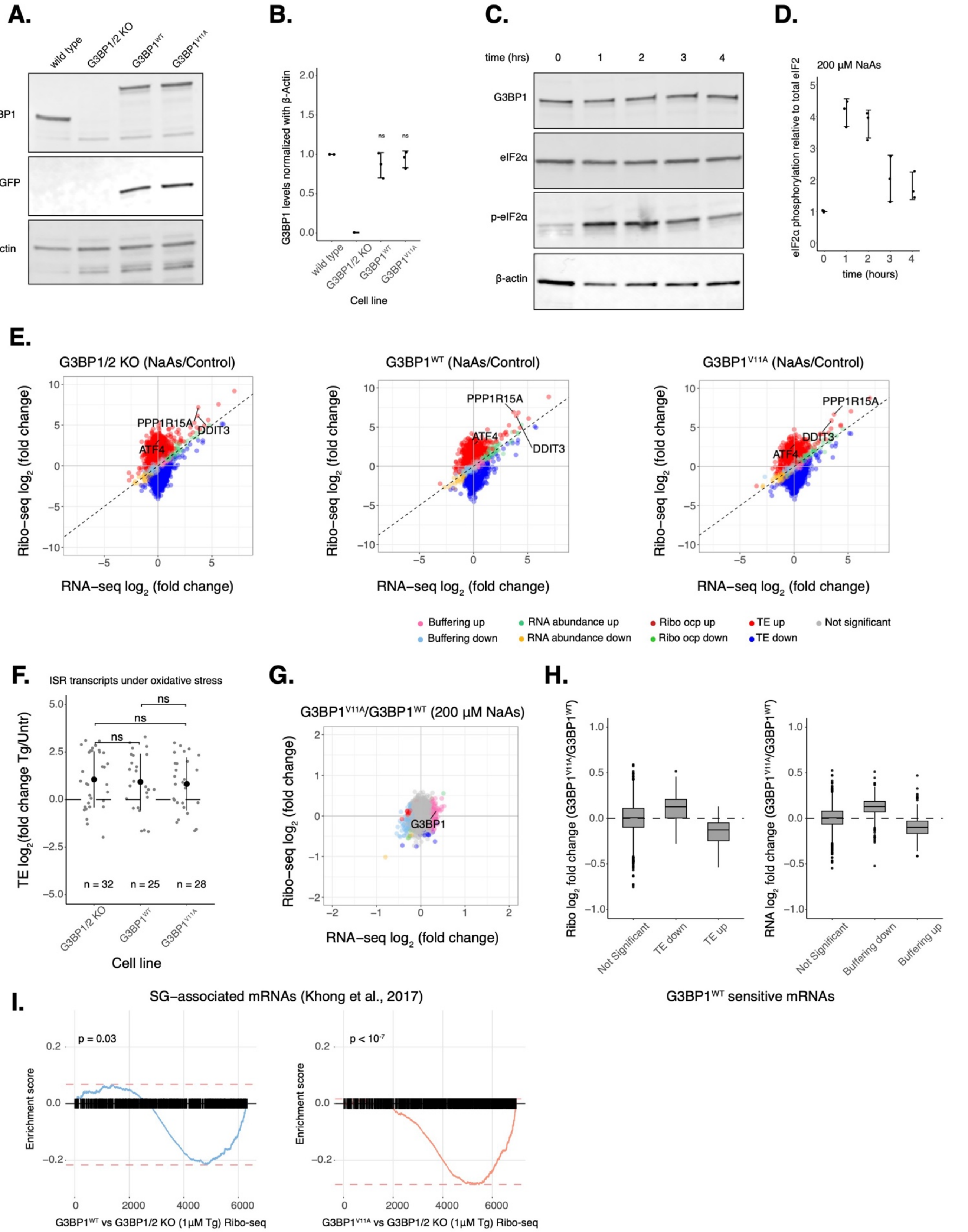

**Figure S6: ISR activation is not affected by G3BP1 condensation under NaAs**. **A**. Western blot showing G3BP1 expression in U2OS wild type, G3BP1/2 KO, $G3BP1^{WT}$, and $G3BP1^{V11A}$ cells. **B**. Quantification of G3BP1 levels across cell lines from data shown in panel A. mean ± SD across $N_{replicates}$ = 3. **C**. Western blot showing a time course of eIF2α phosphorylation for G3BP1/2 KO cells expressing $G3BP1^{WT}$ under 200 μM NaAs. **D**. Quantification of eIF2α phosphorylation across time from data shown on panel C. mean ± SD across $N_{replicates}$ = 3. **E**. Differential expression plots for Ribo-seq and total RNA-seq. G3BP1/2 KO cells or cells expressing either transgenic $G3BP1^{WT}$ or $G3BP1^{V11A}$ were compared between NaAs and Control conditions to show induced expression of canonical ISR factors under NaAs. **F**. Averaged ΔTE of ISR factors across cell lines. Significance was calculated relative to G3BP1/2 KO data. **G**. Differential expression plot for Ribo-seq and total RNA-seq of $G3BP1^{V11A}$ vs $G3BP1^{WT}$ under NaAs. **H**. Ribo-seq (left) and RNA-seq (right) LFC from data shown on panel G. of $G3BP1^{WT}$ sensitive genes identified on Fig. 3D. **I**. GSEA for SG-associated mRNAs overlapping with differentially translated gene sets from $G3BP1^{WT}$ (left) and $G3BP1^{V11A}$ Ribo-seq profiles.

#### Supplemental Figure 7

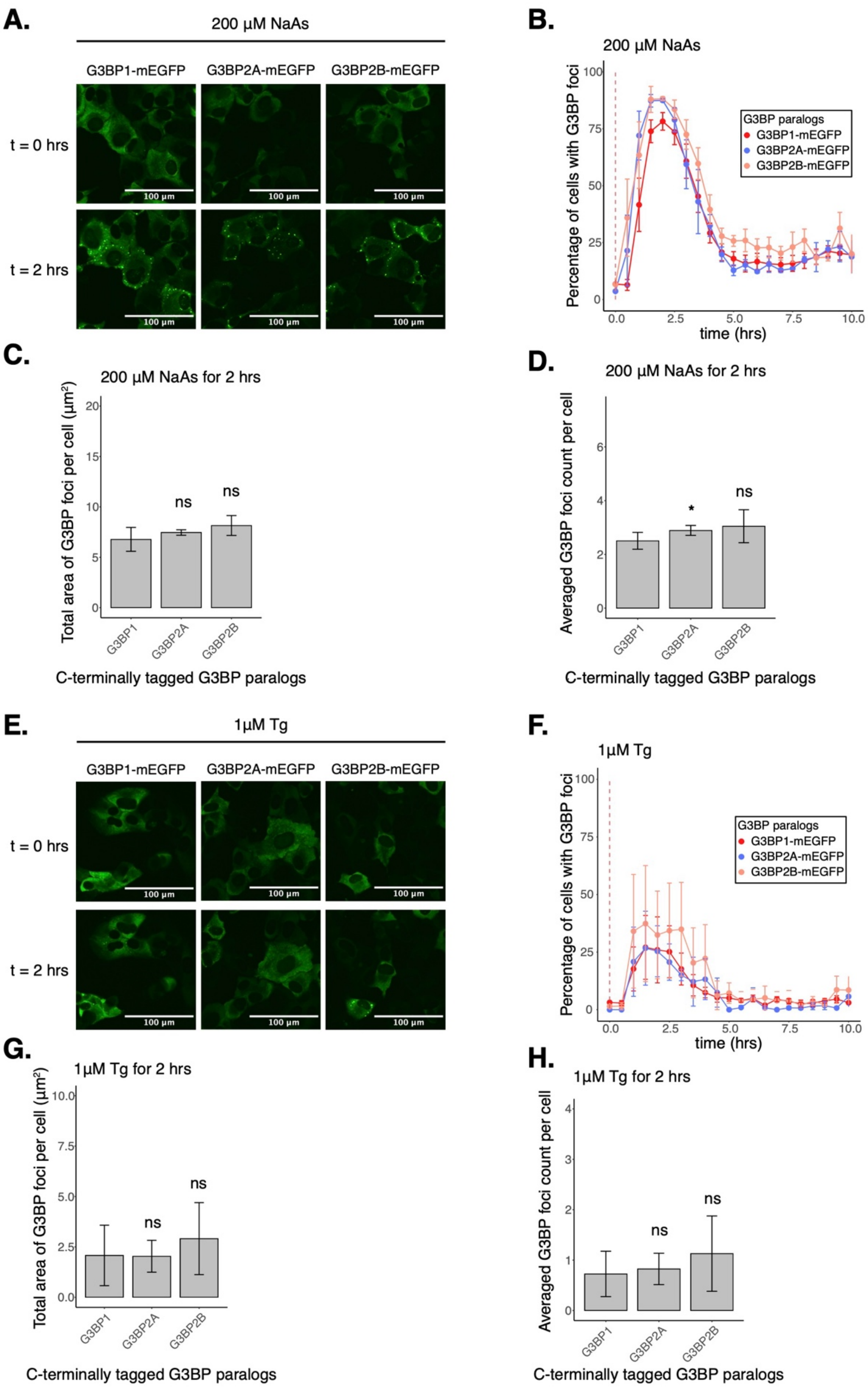

**Figure S7: Condensation of G3BP1/2 paralogs during the ISR**. **A**. Images of cells expressing G3BP-mEGFP paralogs at t = 0 hr and t = 2 hrs post-treatment with 200 μM NaAs. **B**. Percentage of cells with G3BP foci. Vertical red dashed line shows when NaAs was added to cells. **C**. Total area of G3BP foci per cell at 2 hours under NaAs. **D**. G3BP foci count per cell at 2 hours under NaAs. **E**. Images of cells expressing G3BP-mEGFP paralogs at t = 0 hr and t = 2 hrs post-treatment with 1 μM Tg. **F**. Percentage of cells with G3BP foci. Vertical red dashed line shows when Tg was added to cells. **G**. Total area of G3BP foci per cell at 2 hours under Tg. **H**. G3BP foci count per cell at 2 hours under Tg. Plots **B-D** and **F-H** are showing mean ± SEM across $N_{replicates} \geq 3$. P-values were calculated based on whole cell populations ($n_{cells} \geq 100$ per replicate) relative to G3BP1. * $p < 0.05$.

#### Supplemental Figure 8

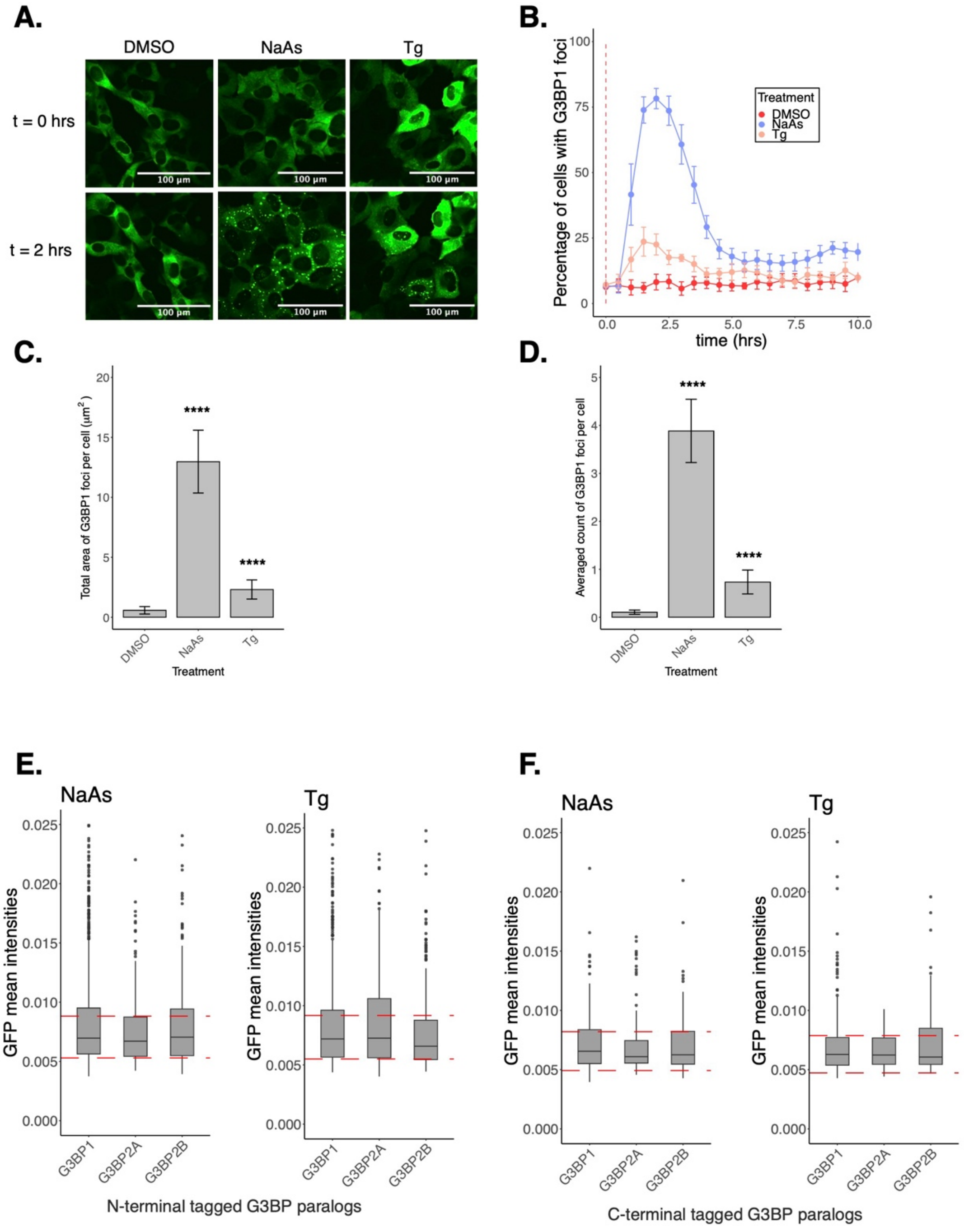


868

**Figure S8: G3BP1 condenses differently across NaAs and Tg stress**. **A**. Images of cells expressing mEGFP-G3BP1$^{WT}$ at t = 0 hr and t = 2 hrs post-treatment with DMSO, 200 μM NaAs, and 1 μM Tg. **B**. Percentage of cells with G3BP1 foci. Vertical red dashed line shows when treatments were applied to cells. **C**. Total area of G3BP1 foci per cell at 2 hours under treatment. **D**. G3BP1 foci count per cell at 2 hours under treatment. Plots **B-D** are showing mean ± SEM across $N_{replicates} \geq 3$. P-values were calculated based on whole cell populations ($n_{cells} \geq 100$ per replicate) relative to DMSO. **** $p < 0.0001$. **E**. Cytoplasmic GFP intensities as a proxy for G3BP1/2 expression across single cells between N-terminal tagged paralogs pre-treated with NaAs and Tg. **F**. Cytoplasmic GFP intensities as a proxy for G3BP1/2 expression across single cells between C-terminal tagged paralogs pre-treated with NaAs and Tg. Horizontal dashed red lines represent ± 25% from G3BP1 median cytoplasmic GFP intensity.

#### Supplemental Figure 9

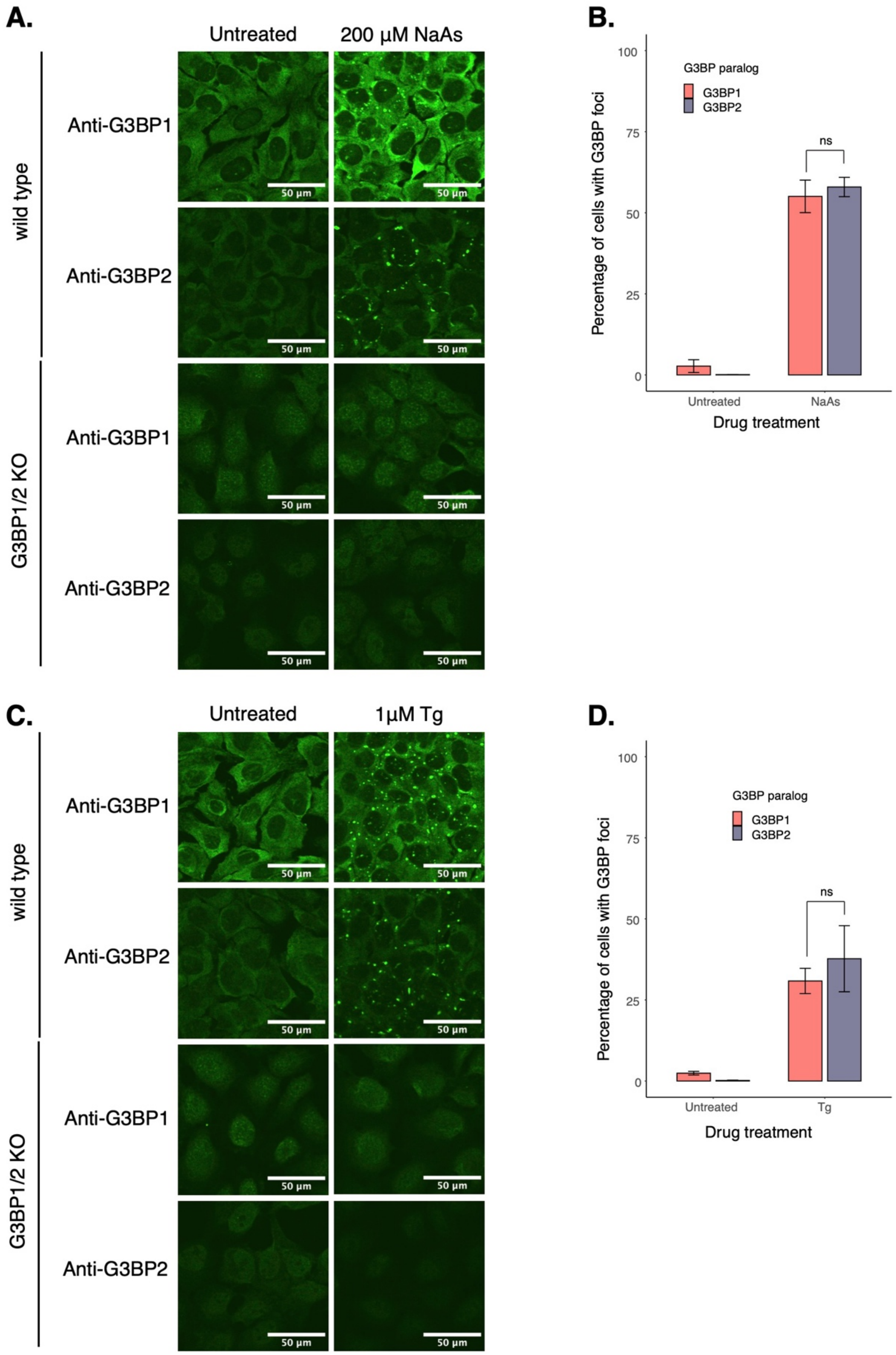

**Figure S9: Endogenous G3BPs condense similarly during the ISR**. **A**. SGs stained against G3BP1 and G3BP2 by IF. Images are showing U-2OS wild type cells and G3BP1/2 KO cells under water or 200 μM NaAs for 2 hours. **B**. Percentage of cells with G3BP1/2 foci from data shown in panel A. **C**. SGs stained against G3BP1 and G3BP2 by IF. Images are showing U-2OS wild type cells and G3BP1/2 KO cells under DMSO or 1 μM Tg for 2 hours. **D**. Percentage of cells with G3BP1/2 foci from data shown in panel C. Plots **B** & **D** are showing mean ± SEM across $N_{replicates}$ = 3.

#### Supplemental Figure 10

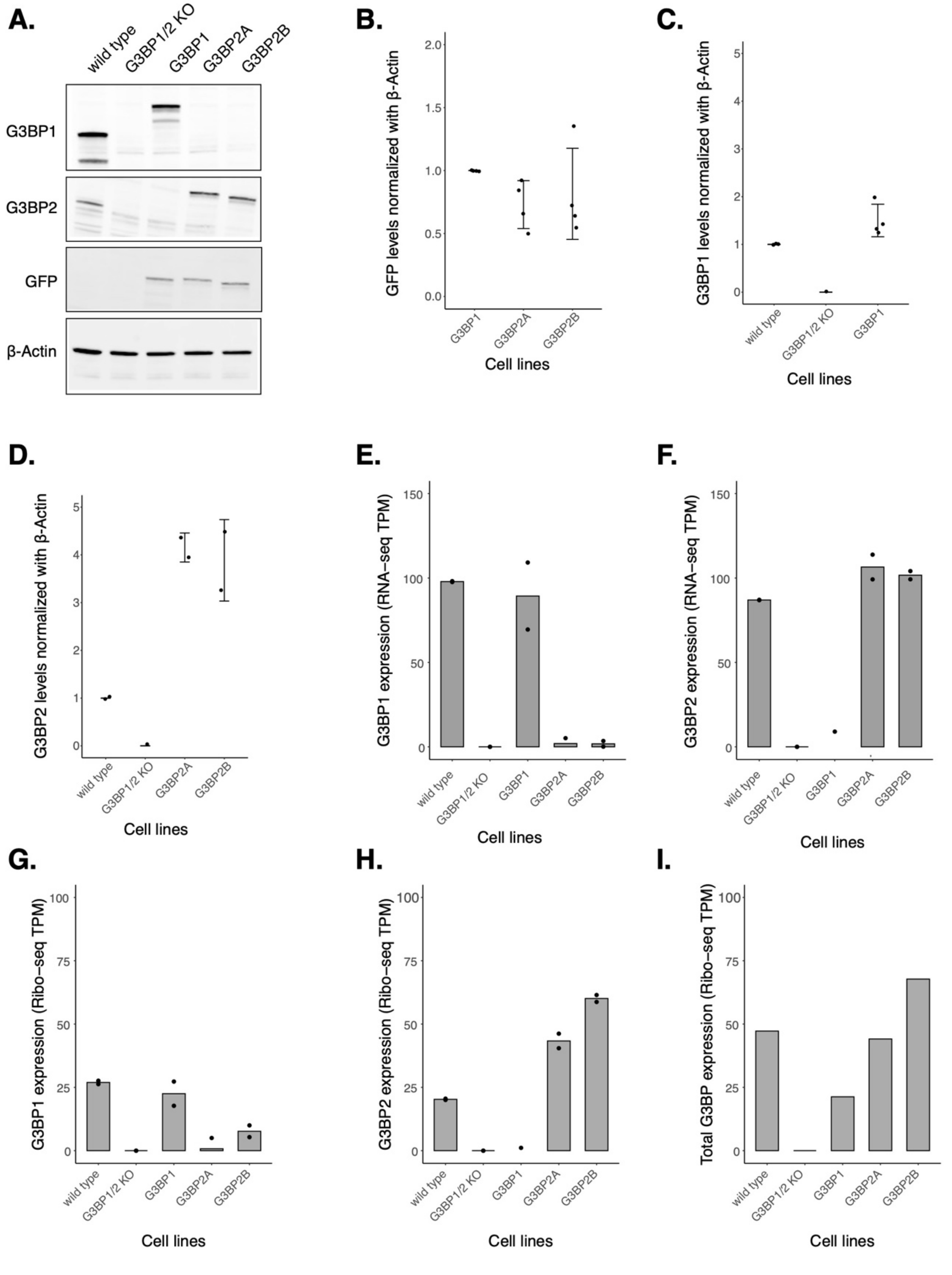


886

**Figure S10: Comparing G3BP1/2 expression across transgenic cell lines**. **A**. Western blot showing expression of G3BPs and GFP across cell lines. **B**. Quantification of GFP levels across transgenic cell lines from data shown on panel A. **C**. Quantification of G3BP1 levels from data shown on panel A. **D**. Quantification of G3BP2 levels from data shown on panel A. For plots **B-D**, mean ± SD. **E**. RNA-seq TPMs of G3BP1 gene across cell lines. **F**. RNA-seq TPMs of G3BP2 gene across cell lines. **G**. Ribo-seq TPMs of G3BP1 gene across cell lines. **H**. Ribo-seq TPMs of G3BP2 gene across cell lines. **I**. Ribo-seq TPMs of total G3BP expression across cell lines.

### Supplemental Figure 11

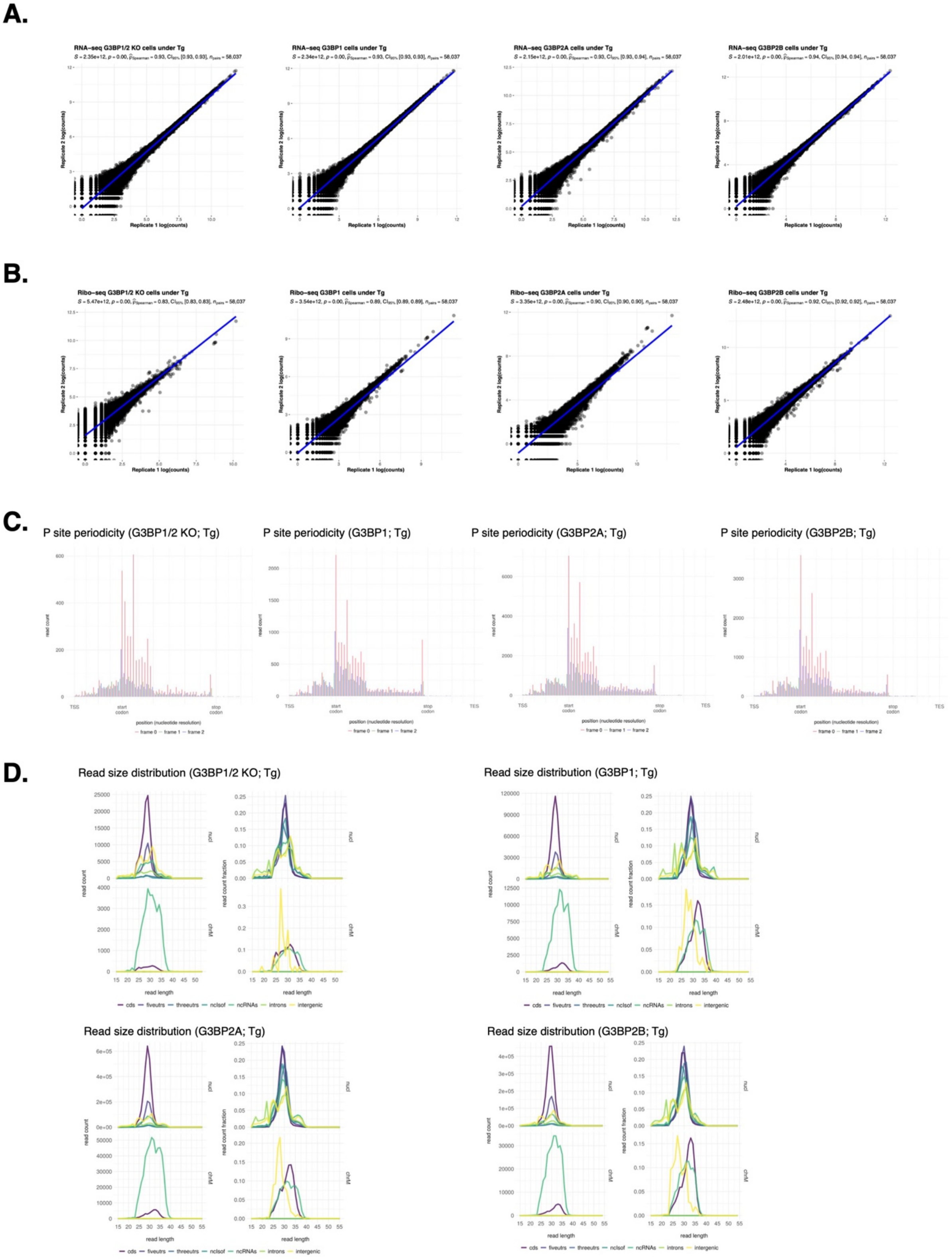

**Figure S11: Sequencing QC data for G3BP1/2 KO, G3BP1, G3BP2A, and G3BP2B profiles**. **A**. Spearman correlation of G3BP1/2 KO (left), G3BP1 (center-left), G3BP2A (center-right), and G3BP2B (left) RNA-seq sample replicates under Tg. **B**. Spearman correlation of G3BP1/2 KO (left), G3BP1 (center-left), G3BP2A (center-right), and G3BP2B (left) Ribo-seq sample replicates under Tg. **C**. P site three nucleotide periodicity for Ribo-seq reads of G3BP1/2 KO (left), G3BP1 (center-left), G3BP2A (center-right), and G3BP2B (left). **D**. Read length distributions of different mRNA species captured by Ribo-seq for G3BP1/2 KO (upper-left), G3BP1 (upper-right), G3BP2A (bottom-left), and G3BP2B (bottom-right)

#### Supplemental Figure 12

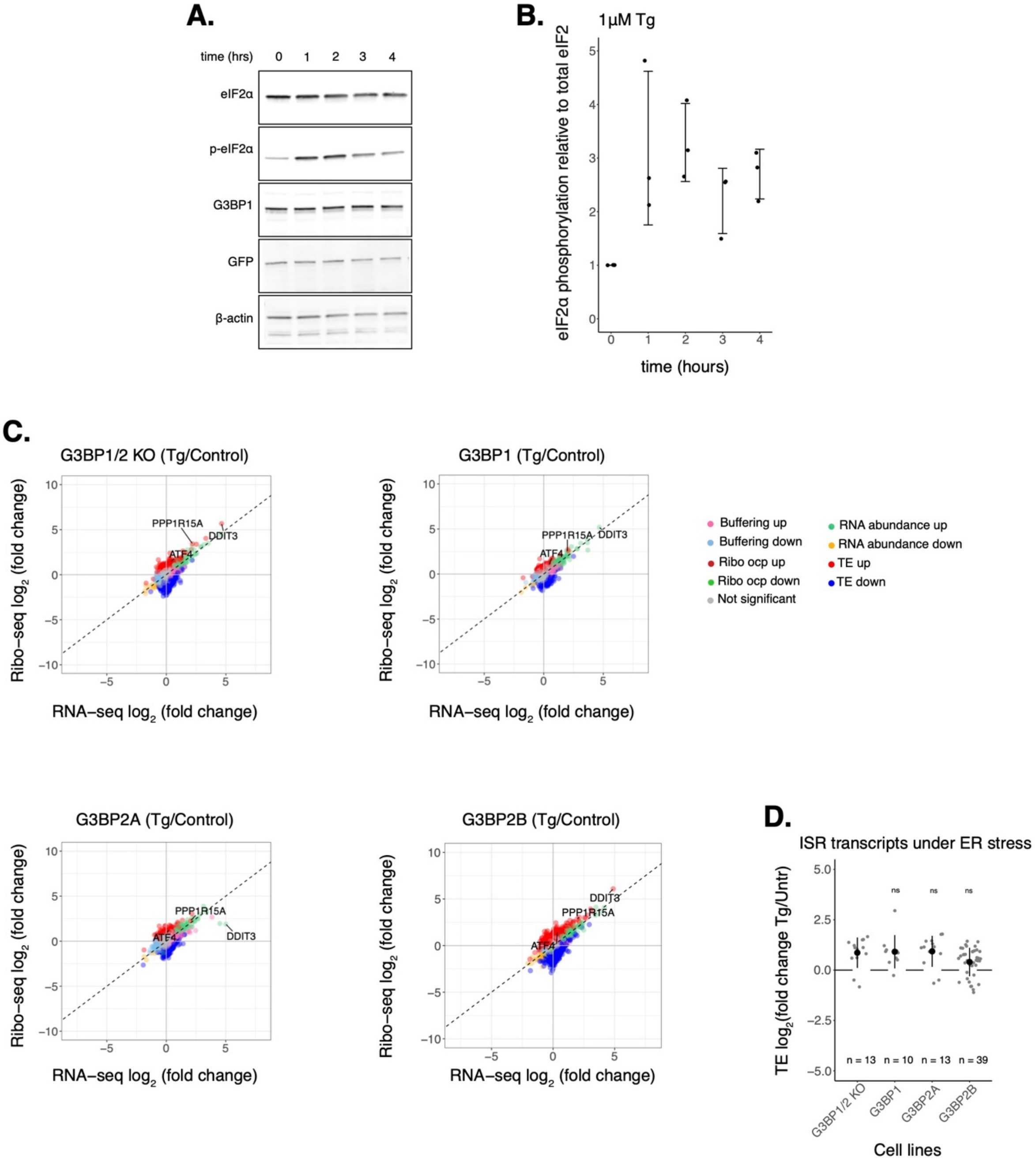

**Figure S12: ISR activation is not affected by G3BP1/2 paralogs under Tg**. **A**. Western blot showing a time course of eIF2α phosphorylation for G3BP1/2 KO cells expressing G3BP1 under 1 μM Tg. **B**. Quantification of eIF2α phosphorylation across time from data shown on panel A. mean ± SD across $N_{replicates} = 3$. **C**. Differential expression plots for Ribo-seq and total RNA-seq. G3BP1/2 KO cells or cells expressing either transgenic G3BP1, G3BP2A, or G3BP2B were compared between Tg and control conditions to show induced expression of canonical ISR factors under Tg. **D**. Averaged ΔTE of ISR factors across cell lines. Significance was calculated relative to G3BP1/2 KO data.

#### Supplemental Figure 13

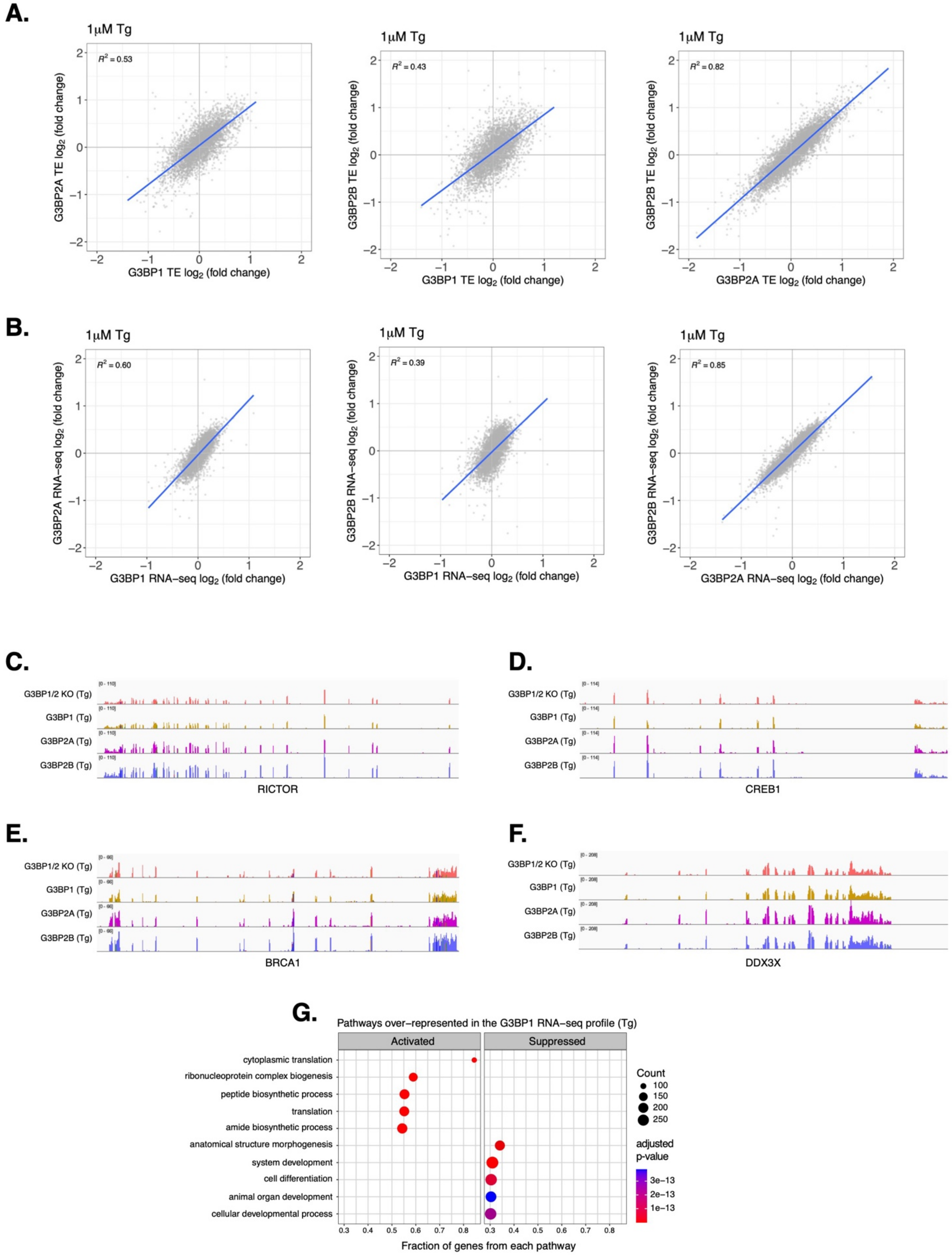

**Figure S13: G3BP1/2 paralogs affect expression of mRNAs differently**. **A**. ΔTE LFC correlations between G3BP1 and G3BP2A (left), G3BP1 and G3BP2B (middle), G3BP2A and G3BP2B (right) under Tg. **B**. RNA-seq LFC correlations between G3BP1 and G3BP2A (left), G3BP1 and G3BP2B (middle), G3BP2A and G3BP2B (right) under Tg. **C-F**. RNA-seq coverage tracks of RICTOR, CREB1, BRCA1, and DDX3X of G3BP1/2 KO, G3BP1, G3BP2A, and G3BP2B expressing cells under Tg. **G**. GSEA identifying activated and suppressed pathways by G3BP1 on the differentially expressed gene sets from RNA-seq under Tg.

**Supplemental Figure 14**

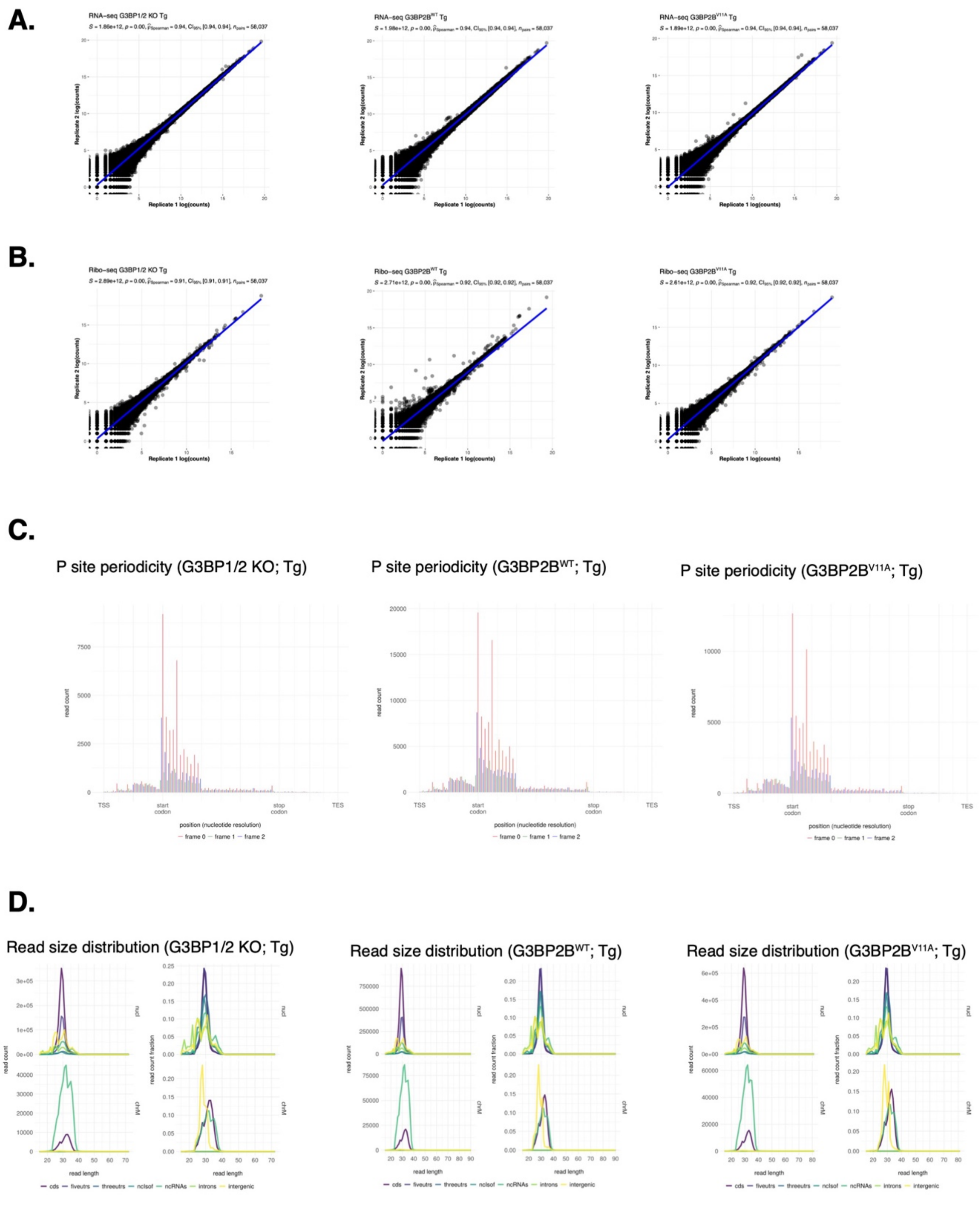


917

**Figure S14: Sequencing QC data for G3BP1/2 KO, G3BP2B$^{WT}$, and G3BP2B$^{V11A}$ profiles**. **A**. Spearman correlation of G3BP1/2 KO (left), G3BP2B$^{WT}$ (middle), and G3BP2B$^{V11A}$ (right) RNA-seq sample replicates under Tg. **B**. Spearman correlation of G3BP1/2 KO (left), G3BP2B$^{WT}$ (middle), and G3BP2B$^{V11A}$ (right) Ribo-seq sample replicates under Tg. **C**. P site three nucleotide periodicity for Ribo-seq reads of G3BP1/2 KO (left), G3BP2B$^{WT}$ (middle), and G3BP2B$^{V11A}$ (right). **D**. Read length distributions of different mRNA species captured by Ribo-seq for G3BP1/2 KO (left), G3BP2B$^{WT}$ (middle), and G3BP2B$^{V11A}$ (right).

#### Supplemental Figure 15

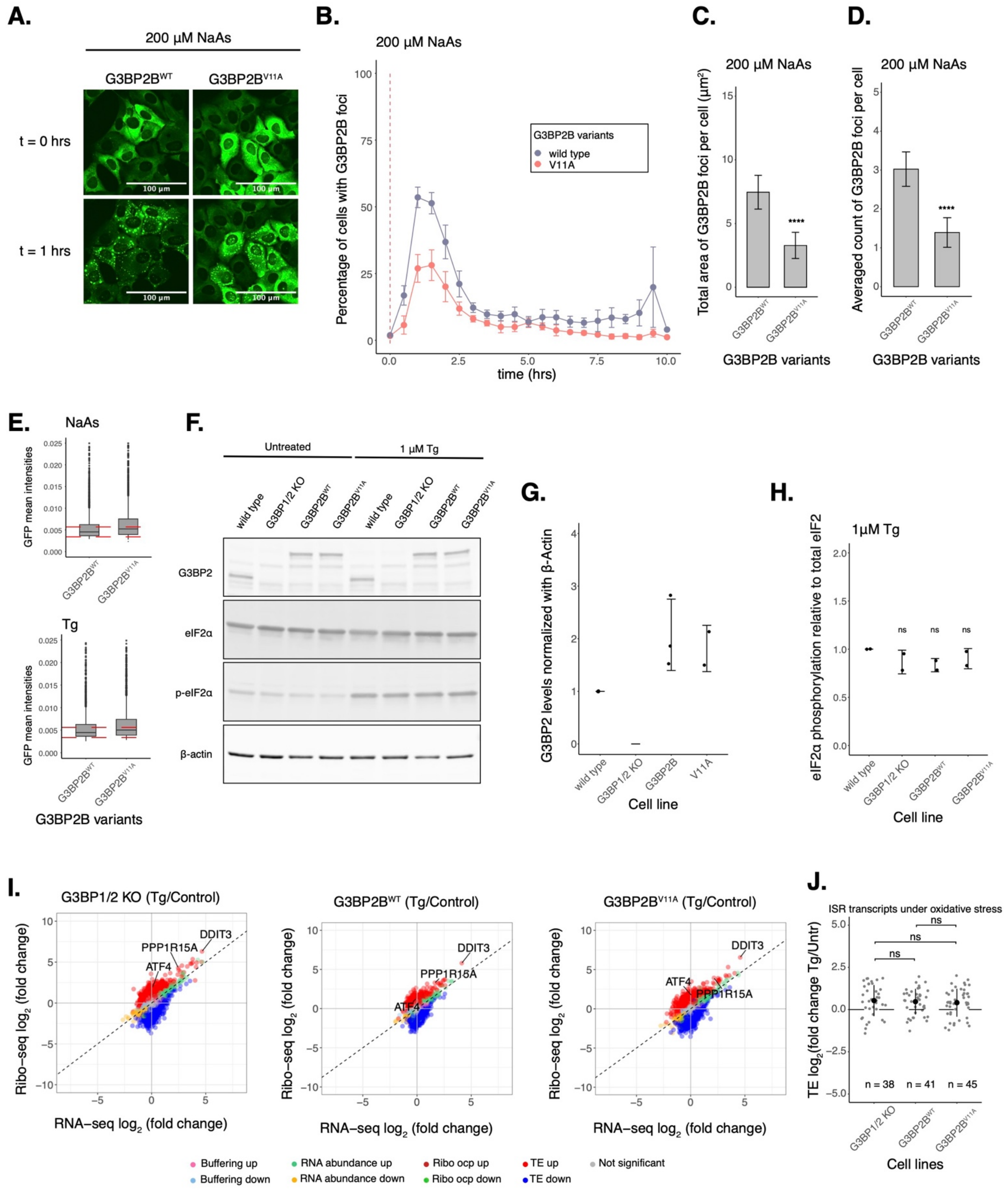

**Figure S15: ISR activation is not affected by G3BP2B condensation under Tg**. **A**. Images of cells expressing mEGFP-G3BP2B variants at t = 0 hr and t = 1 hrs post-treatment with 200 μM NaAs. **B**. Percentage of cells with G3BP2B foci. Vertical red dashed line shows when NaAs was added to cells. **C**. Total area of G3BP2B foci per cell at 1 hour under NaAs. **D**. G3BP2B foci count per cell at 1 hour under NaAs. Plots **B-D** are showing mean ± SEM across $N_{replicates} \geq 3$. P-values were calculated based on whole cell populations ($n_{cells} \geq 100$ per replicate). **** $p < 0.0001$. **E**. Cytoplasmic GFP intensities as a proxy for G3BP2B variant expression across single cells pre-treated with NaAs (upper) and Tg (lower). Horizontal dashed red lines represent ± 25% from G3BP2B median cytoplasmic GFP intensity. **F**. Western blot showing G3BP2 expression and eIF2α phosphorylation in U2OS wild type, G3BP1/2 KO, $G3BP2B^{WT}$, and $G3BP2B^{V11A}$ cells in both DMSO and Tg treated conditions. **G**. Quantification of G3BP2 levels across cell lines from data shown in panel F. mean ± SD across $N_{replicates} = 3$. **H**. Quantification of eIF2α phosphorylation across cell lines from data shown in panel F. mean ± SD across $N_{replicates} = 3$. Significance was calculated relative to wild type cells. **I**. Differential expression plots for Ribo-seq and total RNA-seq. G3BP1/2 KO cells or cells expressing either transgenic $G3BP2B^{WT}$ or $G3BP2B^{V11A}$ were compared between Tg and control conditions to show induced expression of canonical ISR factors under Tg. **J**. Averaged ΔTE of ISR factors across cell lines. Significance was calculated relative to G3BP1/2 KO data.

### Supplemental Figure 16

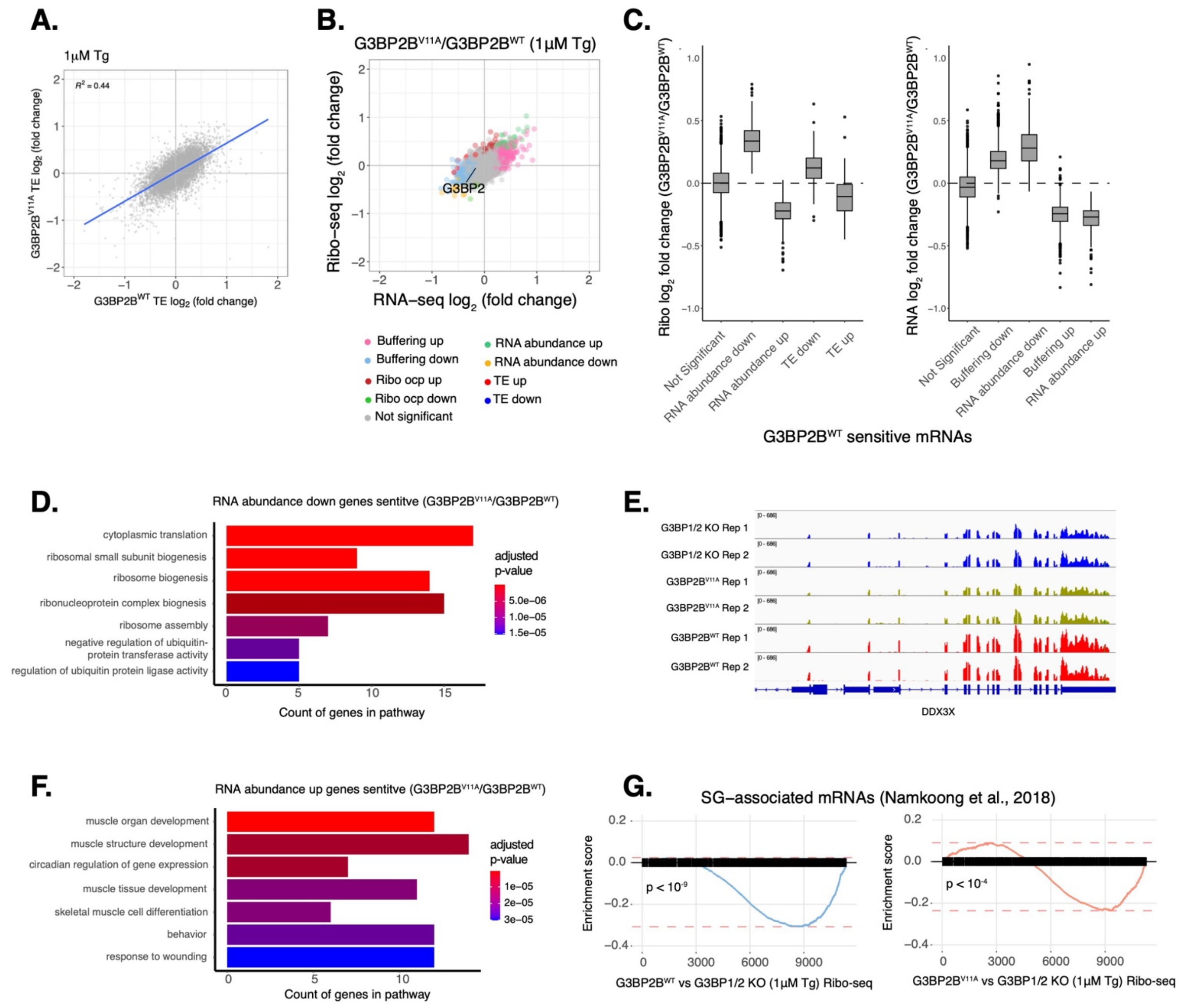

**Figure S16: G3BP2B impacts the expression of select mRNAs under Tg**. **A**. ΔTE LFC correlations between G3BP2B$^{WT}$ and G3BP2B$^{V11A}$ under Tg. **B**. Differential expression plot for Ribo-seq and total RNA-seq of G3BP2B$^{V11A}$ vs G3BP2B$^{WT}$ under Tg. **C**. Ribo-seq (left) and RNA-seq (right) LFC from data shown on panel B. of G3BP2B$^{WT}$ sensitive genes identified on Fig. 6E. **D**. GO of RNA abundance down genes identified in data of panel B. **E**. RNA-seq coverage tracks of DDX3X of G3BP1/2 KO, G3BP2B$^{WT}$, and G3BP2B$^{V11A}$ expressing cells under Tg. **F**. GO of RNA abundance up genes identified in data of panel B. **G**. GSEA for SG-associated mRNAs overlapping with differentially translated gene sets from G3BP2B$^{WT}$ (left) and G3BP2B$^{V11A}$ (right) Ribo-seq profiles.
